# Spatial and molecular anatomy of germ layers in the gastrulating Cynomolgus monkey embryo

**DOI:** 10.1101/2022.01.26.474719

**Authors:** Guizhong Cui, Su Feng, Yaping Yan, Li Wang, Xiechao He, Xi Li, Yanchao Duan, Jun Chen, Patrick P.L. Tam, Ke Tang, Ping Zheng, Wei Si, Naihe Jing, Guangdun Peng

## Abstract

During mammalian embryogenesis, spatial regulation of gene expression and cell signaling are functionally coupled with lineage specification, patterning of tissue progenitors and germ layer morphogenesis. While the mouse model has been instrumental for our understanding of mammalian development, comparatively little is known about human and non-human primate gastrulation due to the restriction of both technical and ethical issues. Here, we present a morphological and molecular survey of spatiotemporal dynamics of cell types populating the non-human primate embryos during gastrulation. We performed serial sections of Cynomolgus monkeys (*Macaca fascicularis*) gastrulating embryos at 1-day temporal resolution from E17 to E21, and reconstructed three-dimensional digital models based on high-resolution anatomical atlas that revealed the dynamic changes in the geography of the mesoderm and primitive streaks. Spatial transcriptomics identified unique gene profiles that correspond to distinct germ layers and cross-species spatiotemporal transcriptome analysis revealed a developmental coordinate of germ layer segregation between mouse and primate. Furthermore, we identified species-specific transcription programs during gastrulation. These results offer important insights into evolutionarily conserved and divergent processes during mammalian gastrulation.

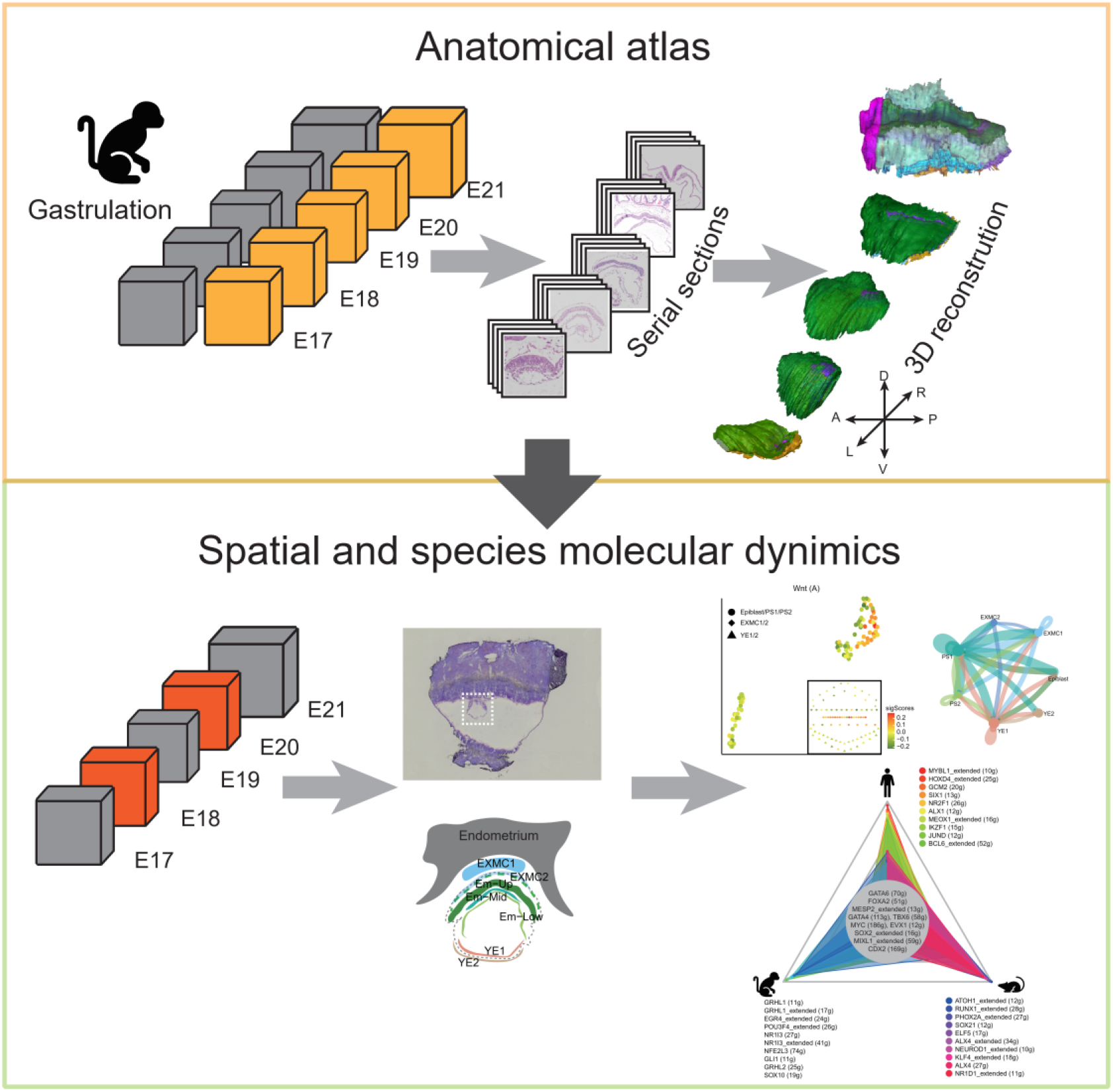

**Highlight:**

- A high-resolution anatomical atlas of Cynomolgus gastrulation embryos
- Created a three-dimensional digital template from serial sections of five developmental stages
- A two-dimensional spatiotemporal transcriptome of the germ layers of gastrulating embryos
- Cross-species comparison infers conservation of functional attributes of regulome and signaling activity in germ layer formation

## Introduction

Gastrulation is fundamental for mammalian embryogenesis at which the definitive germ layers are established and the embryonic cell lineages are allocated to the basic body plan. During germ layer formation, cells undergo ordered changes in shape and fate under a spatiotemporally coordinated regulation. The anatomy of gastrulation has been studied in a variety of mammal species, including the mouse, human and nonhuman primate (Enders et al., 1986; Grobstein, 1985; Mittnenzweig et al., 2021; Moore et al., 1985; Nakamura et al., 2016; Pfister et al., 2007; Snow, 1977; Tarara et al., 1987; Tyser et al., 2020; Yang et al., 2021). In human embryos, the embryonic period is divided into 23 distinct morphological stages known as Carnegie stages (CS) (O’Rahilly and Muller, 2010). At the early CS6 stage (approximately 14 days post-fertilization, Embryonic day 14, E14), the posterior side of the pluripotent epiblast cells start gastrulating to generate three germ layer cells: the ectoderm, mesoderm, and endoderm. However, due to the limitation of the scope of experimental studies of human development up to the onset of gastrulation (Hyun et al., 2021) and the accessibility of embryonic samples, the developmental biology of the gastrulating human embryos is not known in detail.

Previous works have described the gastrulation of non-human primates in the context of morphogenesis and transcriptome, based on a few tissue sections and in vitro models (Hendrickx, 1972; Luckett, 1978; Ma et al., 2019; Nakamura et al., 2016; Niu et al., 2019; Tyser et al., 2020; Yang et al., 2021). However, little is known about the three-dimensional (3D) structure and the molecular landscape, such as the spatiotemporal pattern of gene expression, of gastrulating embryos, both of which are crucial for gaining structural insights of physiology and illustrating fate mapping. Importantly, in light of the kindled interest of in vitro development of human and monkey early post-implantation embryo and the stem-cell based embryo models (Fu et al., 2020; Moris et al., 2020; Niu et al., 2019; Xiang et al., 2019; Zheng et al., 2019), it is imperative to gain the knowledge of in vivo benchmarks of post-implantation development particularly at gastrulation for use as a reference framework for interpreting the outcome of in vitro modeling research. Therefore, high-resolution morphological and spatial transcriptomic analysis of macaque gastrulating embryos is an imperative scientific endeavour.

With the advent of single-cell RNA sequencing, the molecular phenotype of cell types and their developmental trajectory in early mammalian development have been unraveled (Nakamura et al., 2016; Peng et al., 2020b; Pijuan-Sala et al., 2018; Tam and Ho, 2020; Tyser et al., 2020). However, the datasets of single-cell molecular signatures are missing the spatial/positional information of the cells of interest in embryo (Larsson et al., 2021; Peng et al., 2020a). Our previous work showed that spatially resolved transcriptome, not only generates a 3D digital “in situ hybridization” gene expression dataset for mouse gastrulation, also enables the reconstruction of the molecular trajectory of cell populations at gastrulation in time and space (Peng et al., 2019). As the morphogenetic program of immediate post-implantation development of the embryo is different between the mouse and the human, it is far from certain that the rodent-specific developmental mechanism can be extrapolated to the human embryo. It is a widely held view that non-human primate is the more appropriate model for human embryo development (Nakamura et al., 2021). However, systematic profiling of spatial transcriptome in gastrulating monkey embryos has yet to be accomplished, albeit single-cell transcriptome has been documented for Cynomolgus monkey embryo at E6-E17, which did not extend to the gastrulation stages. Here we performed Geo-seq (Chen et al., 2017) to interrogate the molecular architecture and lineage commitment of the Cynomolgus monkey gastrulation embryos. We first established a high-resolution anatomical atlas to describe the key morphogenetic processes and three-dimensional topographical change during three germ layers formation of Cynomolgus monkey embryo. We then established a molecular atlas depicting the spatial pattern of gene expression at two timepoints (E18 and E20) of gastrulation. Finally, we undertook cross-species comparative analysis of spatiotemporal transcriptome data of mouse, Cynomolgus monkey and human embryos to identify molecular programs that may underpin the evolutional convergence, as well as divergence, of the developmental mechanism of gastrulation across these species. Our atlas can be explored via the interactive website.

## Results

### Structural characteristics of gastrulating embryos in non-human primate

Cynomolgus monkey (*Macaca fascicularis*) gastrulating embryos (E17, E18, E19, E20 and E21, equivalent of carnegie stage 6-9) were isolated after in vitro fertilization (IVF) procedure with ethical approval (STAR Methods, Figure 1A-E and S1A-C). The embryos were staged by calculating the gestation days aided by transabdominal ultrasound monitoring. At least two embryo replicates at each developmental stage were studied (Figure S1C) with a very good consistence (Figure 1A-D, S1D-E and Figure S7G). Having the complete representation of embryos of gastrulation stages allow us to build a holistic view of tissue morphogenesis for non-human primates.

**Figure 1.**
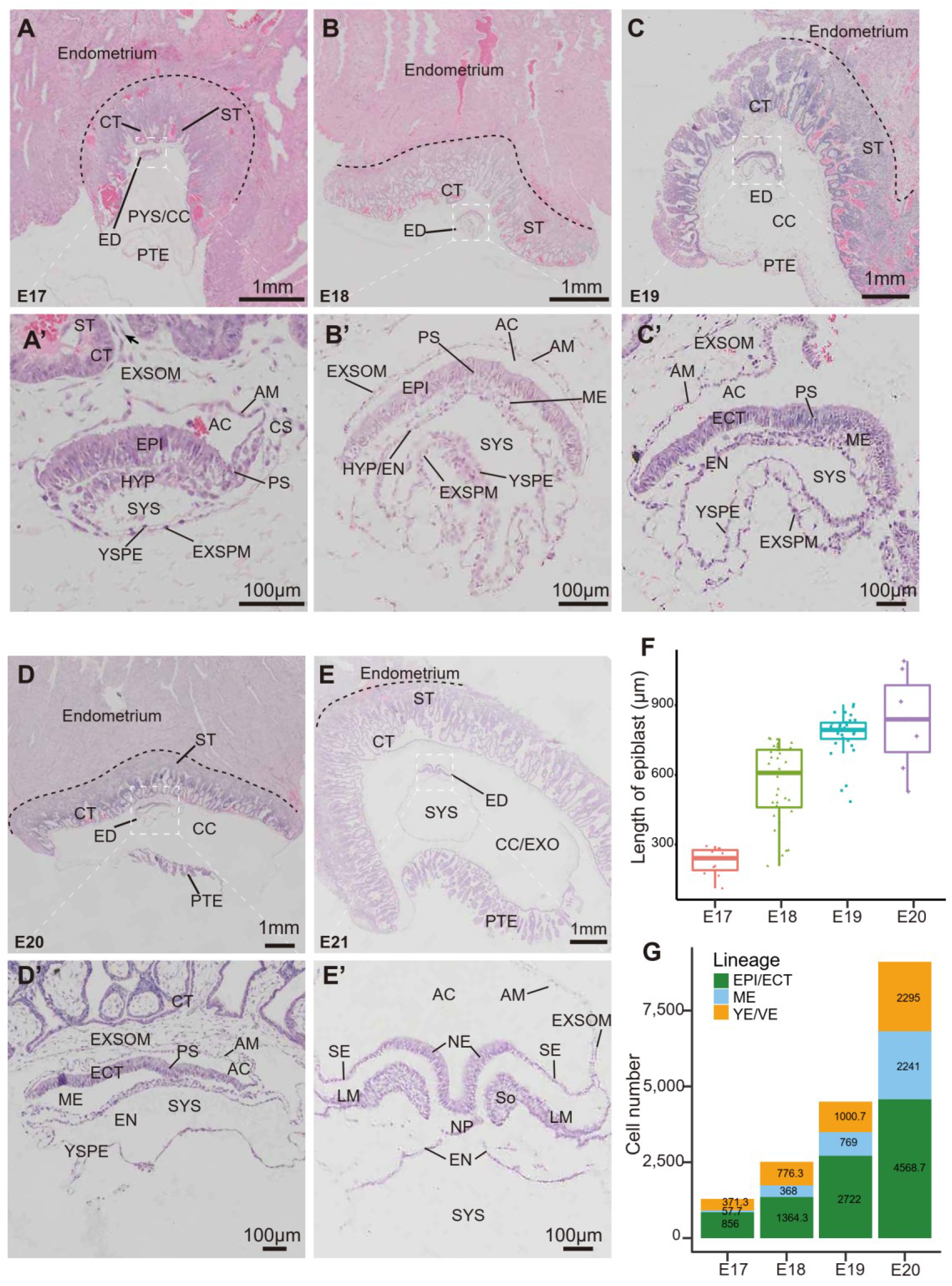
Overview of monkey gastrulation development. A-E, Haematoxylin and eosin staining of the sections of macaque gastrulating embryos at E17 (A and A’), E18 (B and B’), E19 (C and C’), E20 (D and D’) and E21 (E and E’). The image at bottom is a higher magnification of the area boxed on the top. Black arrow points to the extraembryonic mesenchyme has invaded to form the secondary villi. ST, syncytiotrophoblast; CT, cytotrophoblast; PYS, primary yolk sac; CC, chorionic cavity; EXO, exocelom; PTE, parietal trophectoderm; ED, Embryonic disc; EXMC, extra-embryonic mesenchyme cell; AC, amniotic cavity; AM, Amnion; CS, connecting stalk; EPI, epiblast; ECT, ectoderm; PS, primitive streak; ME, mesoderm; HYP, hypoblast; EN, endoderm; YSPE, yolk-sac parietal endoderm; EXSPM, extraembryonic splanchnic mesoderm; EXSOM, extraembryonic somatic mesoderm; SYS, secondary yolk sac. F, The embryo sizes were assessed by the length of sectioned epiblast. G, Bar plot showing total cell numbers in the embryonic germ layers during gastrulation stages. Color represents the germ layer.

Corroborating previous observations on human and rhesus embryos (Enders et al., 1986; Ghimire et al., 2021; Grobstein, 1985; Luckett, 1978; Nakamura et al., 2016), post-implantation Cynomolgus embryos acquired a disc-like configuration and are located on the center of the dome-shaped implantation site which protrudes above the endometrial surface (Figure 1A-E). The extraembryonic mesenchyme has invaded the primary villi to form the secondary villi at E17 (Figure 1A and A’, black arrow). At the tips of secondary villi, cytotrophoblast (CT) cell converged into a thick cytotrophoblastic sheath interspersed with syncytiotrophoblast (ST). Rodent extraembryonic mesoderm originates from the epiblast during gastrulation, whereas the developmental origin of primate extraembryonic mesoderm is hypothesized mainly from the hypoblast-derived primary yolk sac (Bianchi et al., 1993; Ross and Boroviak, 2020). Remarkably, we found that in contrast to mouse, extraembryonic mesoderm of primate embryos was formed before the onset of gastrulation (Figure 1A’). As previously reported, the primate extraembryonic mesoderm constituted the tissues that support the epithelium of the amnion and yolk sac and the chorionic villi (Figure 1A’-E’) (Carlson, 2015). Additionally, the extraembryonic mesoderm splits into two layers, extraembryonic somatic mesoderm (EXSOM) lining the amnion and extraembryonic splanchnic mesoderm (EXSPM) lining the yolk sac (Figure 1A’-E’). These extraembryonic mesoderm cells were subsequently merged with primitive streak (PS)-derived extraembryonic mesoderm to establish the connecting stalk (embryonic stalk), which attaches the whole conceptus to the chorion (Figure 1A’ and Figure S2-6)(Enders and King, 1988). Of note, a unique feature of primate embryos is the formation of secondary yolk sac (SYS) on the hypoblast side the embryonic disc (ED), consisted of an expanding hypoblast (HYP) and squamous yolk sac parietal endoderm (YSPE), gradually expand following gastrulating (Figure 1A’-E’). In the opposite side of ED, amniotic cavity (AC), consisting of a fluid-filled sac surrounding the embryo, was enclosed by low cuboidal epithelium shaped amnion (AM) with epiblast cells.

The formation of the primitive streak (PS) in the midline of the embryonic disc heralds the onset of gastrulation. The epiblast (EPI) was pseudostratified with the properties of typical epithelial cells, and with the elongation of the PS, epiblast cells undergo epithelial-to-mesenchymal transition (EMT) to ingress toward the space between the epiblast and hypoblast, generating mesoderm (ME) wings (Figure 1A’-D’). The anterior-posterior length of epiblast increases from 200μm at early gastrulation (E17) to 900μm at late gastrulation (E20) (Figure 1F and Table S1). The primitive streak spans 20% of the anterior-posterior length of the embryonic disc at E17, increasing to 40% by E20 (Figure S1J). The PS gradually extends to the anterior epiblast, marking a progress of gastrulation (Ghimire et al., 2021; Muller and O’Rahilly, 2004).

### Cell number and cell cycle kinetics of primate gastrulating embryos

A remarkable feature of the development of gastrulating embryos is the regulation of cell growth, especially the mitotic division that results in the increase in cell numbers. In mouse, the embryo that lack normal numbers of cells due to the disruption of cell proliferation or cell death will delay gastrulation until the appropriate number of epiblast cells has been attained (Snow, 1977; Tam and Behringer, 1997). However, the number of cells of the primate embryo during gastrulation remains to be investigated. We obtained a near complete collection of tissue sections, therefore we applied detailed histological analyses on serial sections to determine the kinetics of cell division (STAR Methods, Figure 1G and S1F-I, Table S2). At E17, the early stage of primate gastrulation, the epiblast was about 850 cells, and the hypoblast was approximately 370 cells (Figure 1G and S1K). Subsequently, the number of cells in gastrulating embryos increased rapidly. By the end of gastrulation, the whole embryo was about 9100 cells (4500 cells in the ectoderm, 2200 cells in the mesoderm, and 2300 cells in the endoderm) (Figure 1G and S1K). Analysis of growth rate by determining cell number increase, and by mapping mitotic activity in the germ layers showed that the cell generation time for epithelial ectoderm and endoderm was estimated at about 27 hours (Figure S1K). Notably, the average cell generation time of the mesoderm, perhaps caused by the migration of epiblast and the proliferation of nascent mesoderm, was significantly shortened to 16 hours (Figure S1K). Given that the primitive streak covered about 1/5 of the epiblast at E17 (Figure S1J), the initiation of gastrulation in Cynomolgus monkey embryo was estimated to be approximately at E16 (Nakamura et al., 2021; Nakamura et al., 2016). To assess the threshold number of epiblast cells which is critical for the initiation of gastrulation, the epiblast and hypoblast at starting gastrulation were calculated about 490 and 200 cells, respectively, based on the kinetics of cell proliferation in macaque embryo. Interestingly, the mean cell generation time during primate gastrulation was the half of pre-gastrulation, which is consistent with the pattern found in mouse (Figure S1L) (Snow, 1977). In addition, the length of epiblast progressively increases and the size of the embryo becomes larger (Figure 1F, Table S1). That is slightly different from the in vitro culture system (Niu et al., 2019). Both in vivo and in vitro embryos have similar same size at the early stage of gastrulation, but subsequently, natural embryos become larger while cultured embryos do not change too much in size. This discrepancy may be due to the limitation of the in vitro culture system for complex structural development.

### Cellular anatomy of primate gastrulation embryo

Morphogenetic movements including cell migration and reorganization during gastrulation reshape the embryos. A paramount event is the epithelial to mesenchymal transition (EMT). EMT involves changes of epiblast cells from a proper epithelium with the full array of epithelial characteristics to mesenchymal-shaped mesoderm cells (Massri et al., 2021). During the initiation of mouse gastrulation, the PS, a transient embryonic structure where gastrulation EMT took place, was formed at the posterior of the epiblast. We observed that EMT follows similar patterns in monkey embryos (Figure 2 G-H and G’-H’, white open arrowheads). The epiblast cells were contiguous with amnion cells at the margin of the disc, where there was a gradation in cell size from columnar to low cuboidal spanning two to three cells (Figure 2 E-F and E’-F’, asterisks). Cells that remain in the epiblast are allocated to the ectoderm (ECT) (Ghimire et al., 2021; Shahbazi, 2020). A thick basement membrane lining the epiblast surface was visible in the anterior region (Figure 2 A-C and A’-C’, black arrows), while epiblast cells in the primitive streak were still connected to the rest of the epiblast with the onset of gastrulation (Figure 2 G-H and G’-H’, white open arrowheads). These cells changed their shape from tall column to triangle, lost the epiblast cell–basement membrane interaction, and were squeezed together. During gastrulation, we found that cells ingress through the PS, break down the basement membrane, and invade into the space between the hypoblast and epiblast to form the intraembryonic mesoderm (Figure 2 E-H and E’-H’, black open arrowheads). There were also cells that migrated posterolaterally beyond the embryonic disc border to form extraembryonic mesoderm (Figure 2 F-H and F’-H’, white arrows). Thus, the epiblast undergoes a drastic morphological transition, engaging intra/extraembryonic mesoderm cells into a migratory behavior.

**Figure 2.**
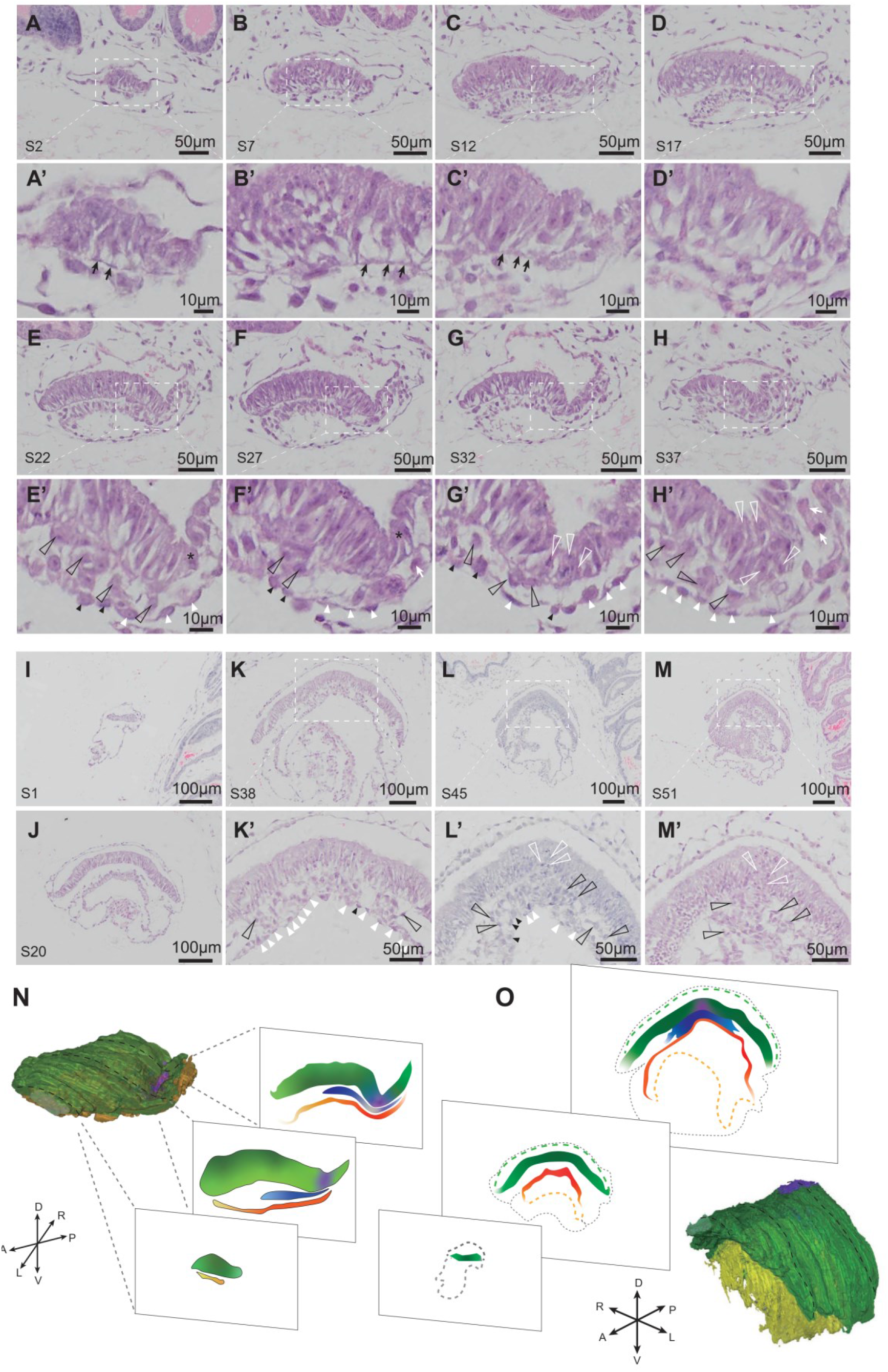
Geography of early gastrulation stages in monkey embryos. A-M, Haematoxylin and eosin staining of the sections of macaque gastrulating embryos at E17 (A-H and A’-H’) and E18 (I-M and K’-M’). The number of the section is shown in the lower left. High-magnification images are from the boxed region. Black arrows point to basement membrane underlying the epiblast, and white arrows point to extraembryonic mesoderm cells. White open arrowheads indicate gastrulating cells in primitive steak, with black open arrowheads indicating mesoderm cells. White closed arrowheads indicate hypoblast cells, with black closed arrowheads indicating definitive endoderm cells that intercalate in the overlying visceral endoderm epithelium. Asterisks mark embryonic/extraembryonic borders. Scale bar as indicated. A, anterior; P, posterior; L, left; R, right; D, dorsal; V, ventral. N-O, Three-dimensional view of the reconstructed embryo highlighting the primitive streak and germ layers with different colors at E17 (N) and E18 (O). The black dash lines indicate the level of the transversal sections.

Notably, some gastrulating cells posterior to the leading edge of the mesoderm wrings were found to undergo mesenchymal epithelial transition (MET), the reverse mechanism of EMT, as they intercalate into the overlying hypoblast (visceral endoderm) epithelium to form the gut endoderm on the surface of the embryo (Figure 2 E’-G’, K’-L’ and Figure 3 C’-E’,K’-M’, black arrowheads). These MET cells, with obvious mesenchymal characteristics, were larger than the pre-existed hypoblast cells but were not in a dominative presence. As previously reported that there were two populations of definitive endoderm cells in human gastrulation embryos by single cell transcriptome analysis (Tyser et al., 2020), this may suggest that the mixing of these two (embryonic and extraembryonic) endodermal populations collectively comprises the definitive gut endoderm, as they do in the mouse embryo (Viotti et al., 2014).

**Figure 3.**
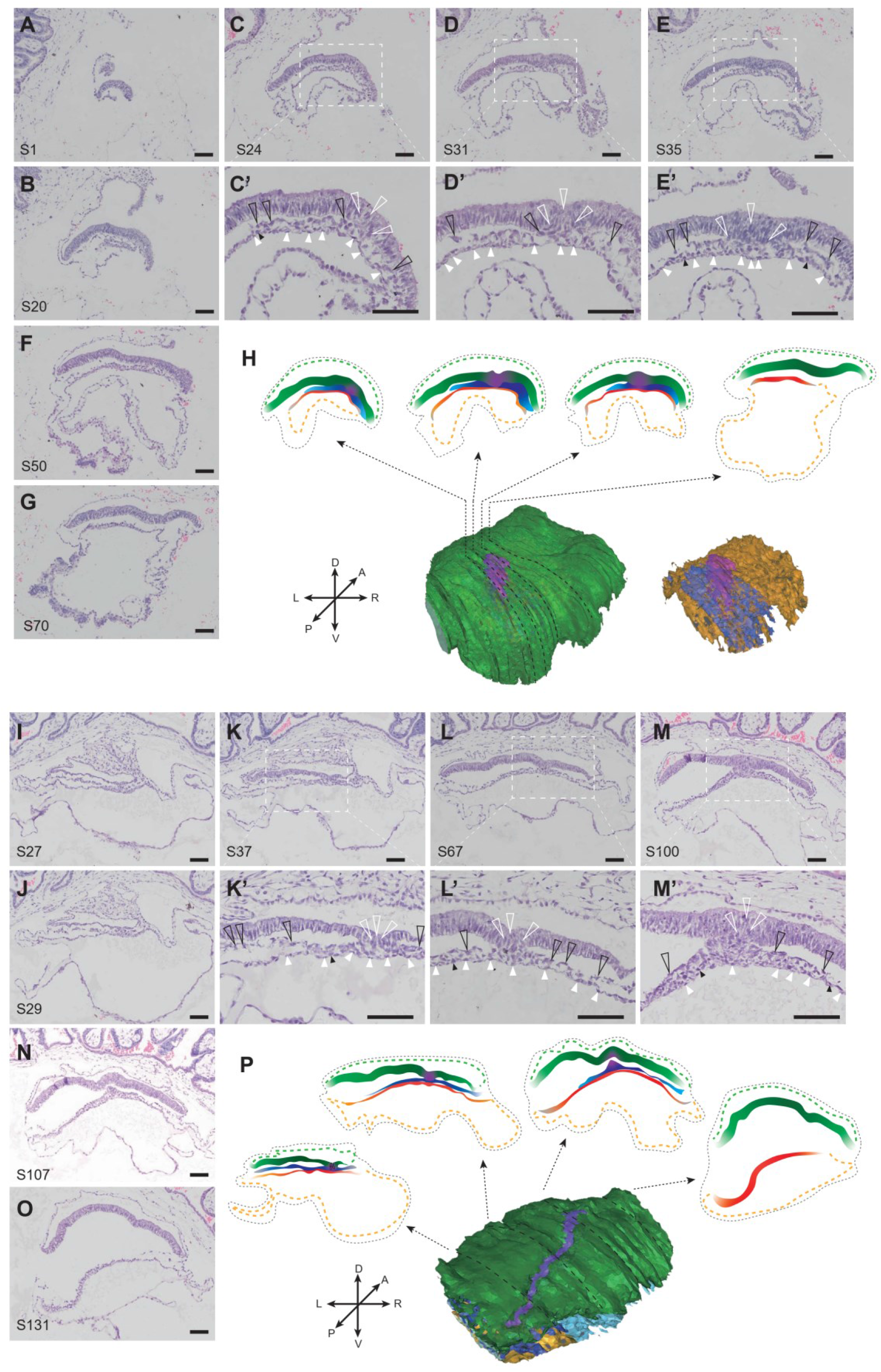
Geography of late gastrulation stages in monkey embryos. A-G, Haematoxylin and eosin staining of the sections of macaque gastrulating embryos at E19. The number of the section is shown in the lower left. High-magnification images are indicative of the boxed region. White open arrowheads indicate gastrulating cells in primitive steak, with black open arrowheads indicating mesoderm cells. White closed arrowheads indicate hypoblast cells, with black closed arrowheads indicating definitive endoderm cells that intercalate in the overlying visceral endoderm epithelium. Scale bar, 100μm. H, Three-dimensional view of the reconstructed embryo highlighting the primitive streak and germ layers with different colors at E19. The black dash lines indicate the level of the transversal sections. I-O, Haematoxylin and eosin staining of the sections of macaque gastrulating embryos at E20. P, Three-dimensional view of the reconstructed embryo at E20. The black dash lines indicate the level of the transversal sections.

Next, to address the intricate morphogenesis of the developing primate embryo, the three-dimensional (3D) anatomical atlas was performed based on serial sections with the collected gastrulation embryos. In line with knowledge from 3D reconstruction of human specimen, we found that the epiblast of Cynomolgus monkey changed from a globe-like into a disc-like shape, with the primitive streak in the midline (Figure 2N-O and Figure 3K, P). Afterwards, the flat disc shape was distorted as the embryonic cavity increases and perfect morphology is difficult to be preserved without perfusion (Figure 3 O, P and S5). At the end of gastrulation, the embryo which begins as a flat sheet of cells, folds to acquire a typical cylindrical shape (Figure 4 and S6). The ectoderm developed into neural ectoderm and surface ectoderm, then neural ectoderm underwent bending to create the neural folds, which converge towards the dorsal midline (Figure 4 A-C and E) (Nikolopoulou et al., 2017). The mesoderm continually developed in the tail bud (Figure 4 D), and divided into the paraxial mesoderm, intermediate mesoderm, and lateral plate mesoderm (Figure 4 C) (Tani et al., 2020). The paraxial mesoderm gives rise to the somite, a notable feature of early organogenesis, mainly through somitogenesis (Lawson and Wilson, 2016). Besides, somite count is a convenient means to stage embryos, and there were 5 or 6 somites visible at E21, corresponding to CS9 of human embryo development (Figure 4 C and E). After gastrulation, the gut endoderm was regionalized along the dorsal-ventral (D-V) and anterior-posterior (A-P) axes into broad foregut, midgut, and hindgut domains. In the monkey embryos, we observed that the foregut pocket invaginated to generate the anterior intestinal portal, while the hindgut pocket at the posterior formed the caudal intestinal portal, a process similar to that in the mouse embryo development (Figure 4 E and S6) (Nowotschin et al., 2019). Together, we reconstructed the histologic detail based on more than 100 serial high-quality sections and provided a digitalized anatomy atlas for the Cynomolgus gastrulation monkeys which can serve as valuable reference for primate embryogenesis.

**Figure 4.**
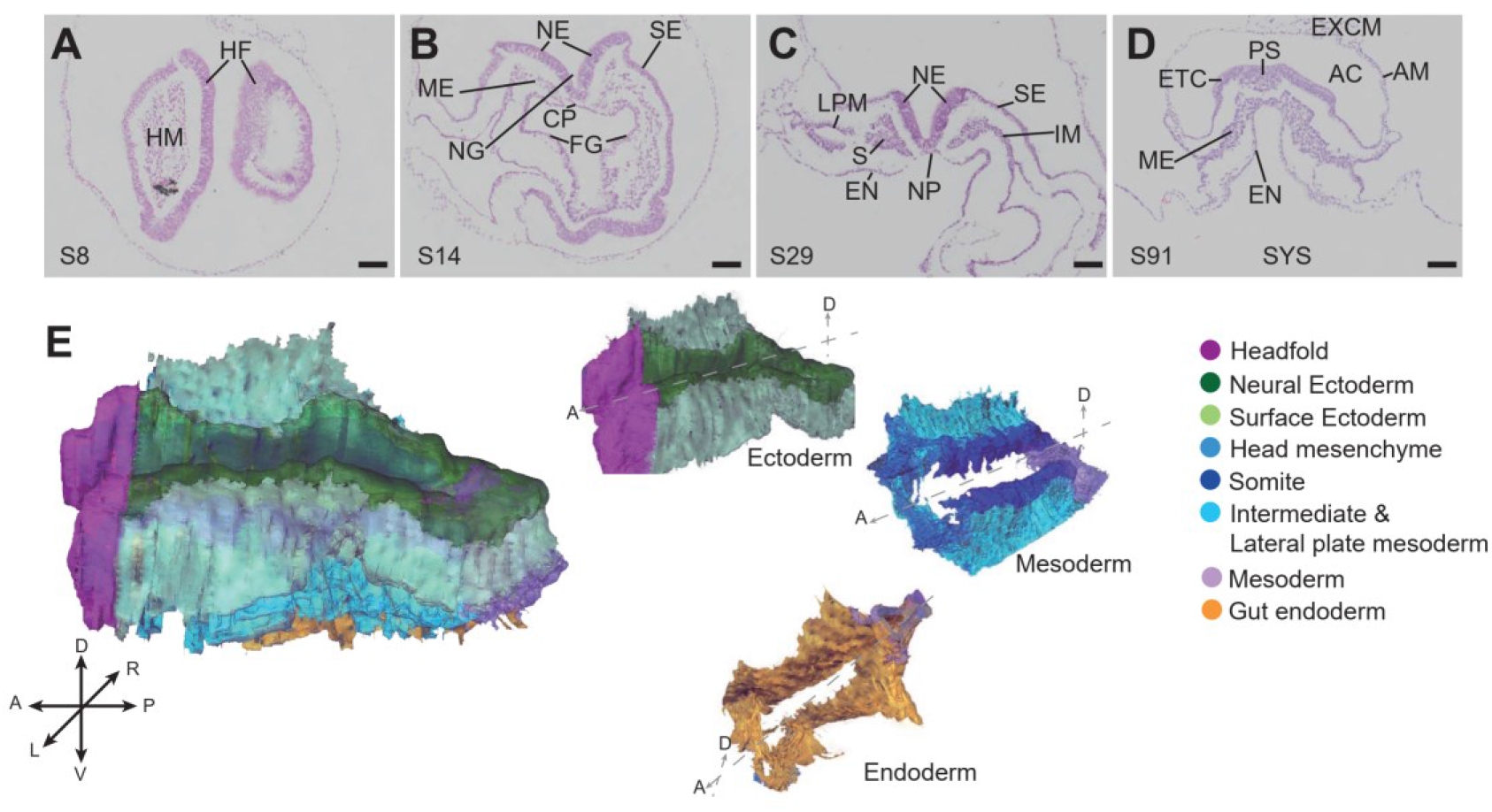
Morphology of primate early organogenesis. A-D, Haematoxylin and eosin staining of the sections of macaque organogenesis embryos at E21. The number of the section is signed in the lower left. HF, headfold; HM, head mesenchyme; FG, foregut; NG, neural groove; SE, surface ectoderm; NE, neural ectoderm; CP, chordal plate; ME, mesoderm; S, somite; NP, notochordal plate; IM, intermediate mesoderm; LPM, lateral plate mesoderm; EN, endoderm; PS, primitive streak; ECT, ectoderm; EXMC, extra-embryonic mesenchyme cell; AC, amniotic cavity; AM, Amnion; SYS, secondary yolk sac. Scale bar, 100μm. E, Three-dimensional view of the reconstructed embryo at early organogenesis stage (E21). A, anterior; P, posterior; L, left; R, right; D, dorsal; V, ventral.

### Molecular architecture of the gastrulating monkey embryo

Previously, we conducted a spatiotemporal transcriptome on the mouse embryo by low-input Geo-seq during gastrulation from E5.5 to E7.5 stages (Peng et al., 2019). To achieve a spatial molecular profiling of primate embryos during gastrulation, we applied Geo-seq to analyze the expression of genes in embryonic and extraembryonic tissues at E18 and E20 (Figure 5A and S7A-C, STAR Methods). Although the tissue morphology of cryo-sections was inferior to paraffin embedding sections, the germ layers were discernable. We divided the embryonic tissues into the upper, middle and lower layer (Em-Up, Em-Mid and Em-Low) according to the dorsal-ventral axis based on the anatomical structure. Furthermore, extra-embryonic mesenchyme 1 and 2 (EXMC1/ EXMC2) and yolk-sac parietal endoderm (YE1) and extraembryonic mesemchyme lining the yolk sac (YE2) were captured from extraembryonic tissues according to their locations in the amniotic cavity and secondary yolk sac (Figure S7A-C). The spatial transcriptome of primate gastrulating embryos provides a high-depth and quality dataset, with ∼15M mapped reads, ∼87% mapping ratio, and 12,000 genes on average (Figure S7F). The consistency between two replicates or developmental stages was more than 0.94, indicating we can rely on these data to probe the molecular architecture of the primate embryo (Figure S7F). We further generated a dot plot to facilitate the planar (top represents dorsal and bottom represents ventral) display of the spatial expression of the combined gastrulating primate embryos, which called STGE-plot (Spatial Transcriptome of Gastrulating Embryos), according to the geography and morphology of qualified samples (n=86) (Figure 5A).

**Figure 5.**
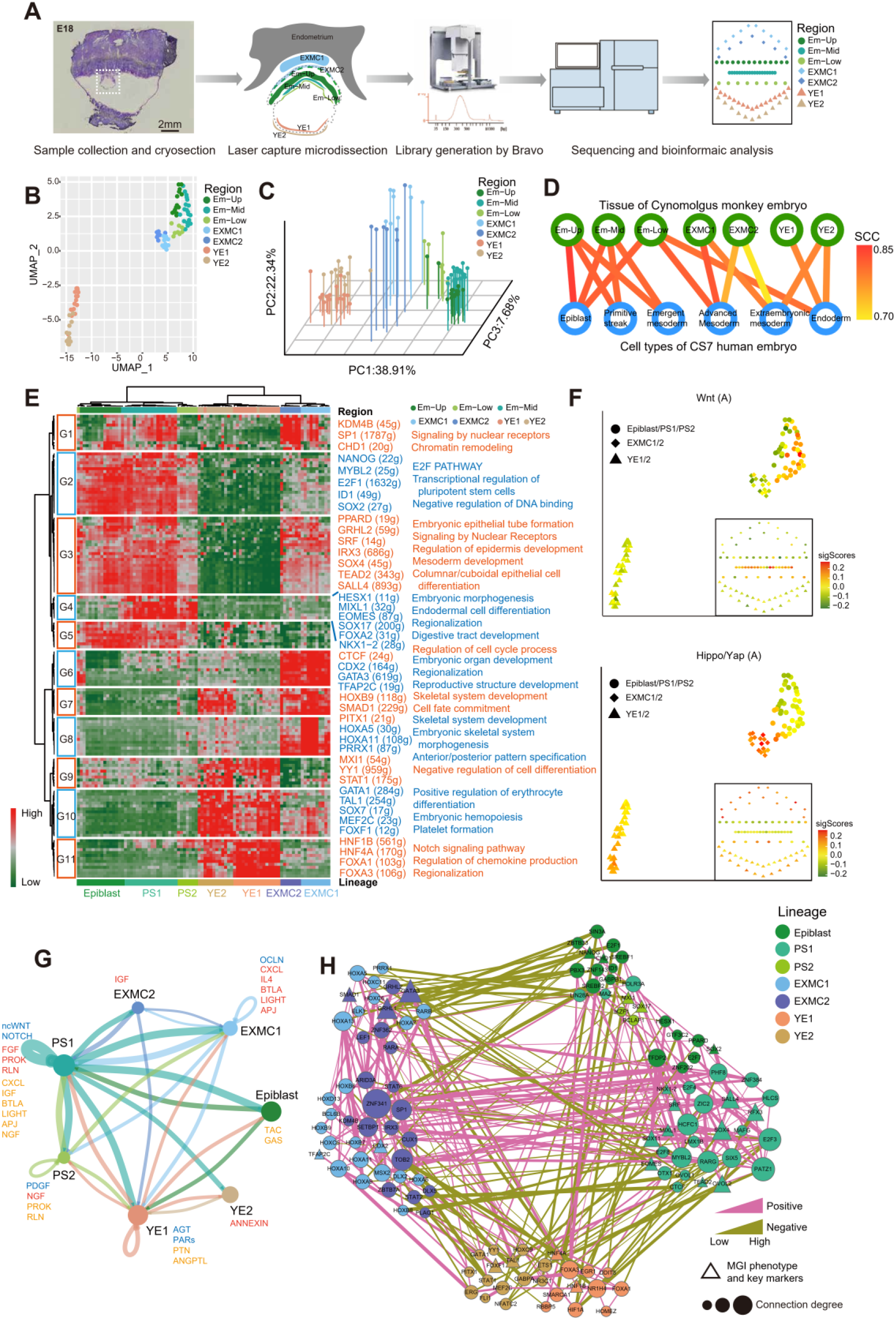
Embryonic and extraembryonic spatial molecular dynamic of gastrulating monkey embryos. A, Schematic overview of spatial transcriptome of monkey embryo by Geo-seq. The top to bottom of the STGE-plot represents the dorsal-ventral axis of the embryo, which are extra-embryonic mesenchyme tissues (EXMC1/2), embryonic disk (Em-Up, Em-Mid and Em-Low) and yolk-sac endoderm (YE1/2), while left-right does not represent craniocaudal axis of embryo. B, Uniform manifold approximation and projection (UMAP) embedding of monkey samples based on high variable genes (n=5,000) colored by spatial location (n=86). C, 3D-PCA plot based on the regulon activity scores of embryonic and extraembryonic samples showing separate spatial domains. D, The Spearman correlation coefficient of cell population between Cynomolgus monkey spatial (red module) and human single cell annotated cell types (blue module) showing the location of specific cell. The color coding is spearman correlation coefficient (SCC). E, The heatmap of specific regulons (n=192) showing 11 regulon groups in tissue samples of gastrulating monkey embryos with listing of examples of regulon transcription factors (numbers of predicted target genes by SCENIC in the brackets) and the enriched Gene Ontology (GO) terms for each regulon group. F, UMAP and STGE-plot showing the activities of the target genes related to the activated states of the Wnt and Hippo–Yap signalling pathway in primate gastrulation. G, The weighted interaction strength between different tissue types by Cellchat. The autocrine and paracrine (outgoing and incoming) signaling pathways were colored by blue, red and orange respectively. Edge width represents the communication probability. H, The co-activation network of tissue specifical regulons based on spearman correlation coefficient (SCC >0.9). The node with different colors represents the highest activity tissue and the edges are the SCC value of two nodes. Violet lines indicate the positive correlation and green lines indicate negative correlation. A wider edge signifies higher correlation. Triangle nodes denote transcription factors with mouse knockout gastrulation phenotype in MGI database. The larger the node, the more connections to other regulons.

Next, we performed principal component analysis (PCA) and uniform manifold approximation and projection (UMAP) of the tissue samples with unsupervised clustering, and the result showed tight clustering of samples by spatial positions (Figure 5B), pointing to a location-dependent germ layer specification. To characterize the functional status of gene regulatory network (GRN), we performed SCENIC (single-cell regulatory network inference and clustering) (Aibar et al., 2017) analysis (STAR Methods). The specifically enriched regulon activity based on transcription factors and their co-expressed target genes was further grouped into 11 modules that are specifically activated in seven spatial cell clusters (Figure 5 C and E). In line with the unsupervised clustering of transcriptome, the regulon-based dimensionality reduction showed that the tissue type and spatial location were clearly delineated on the top three PC axes (Figure 5C). The PC1 axis defined cell types of embryonic and extra-embryonic tissues, and the PC2 and PC3 axis represented the spatial variances. Therefore, the signatures of tissue types and geographical regions of Cynomolgus monkey gastrulating embryos were distinguished by both gene expression and regulatory activity.

To investigate the lineage identities of spatial samples, we incorporated key marker genes, regulon activities, geographical anatomy and cell-type deconvolution to infer spatial molecular organization of primate gastrulation. For cell-type deconvolution in spatial transcriptome, SPOTlight that based on a non-negative matrix factorization (NMF) regression algorithm was used to infer the location of cell types (Elosua-Bayes et al., 2021). We integrated both in vivo single cells (Nakamura et al., 2016) and our spatial transcriptomics data of monkey gastrulating embryo, and demonstrated that gastrulating cells (Gast1, 2a and 2b) were mainly located in middle and lower layer of embryonic disk whereas the post-implantation late epiblast cell (PostL.EPI) account for the largest proportion in the upper layer of embryo (Figure S7D). In line with cell-type deconvolution analysis, the correlation coefficiency of gene expression showed that the upper layer of embryonic tissues was more similar to the epiblast cells, whereas there was a discrete regional property of embryonic and extra-embryonic cell types (Figure S7E). In addition, the spatially captured tissue from gastrulating monkey embryos were consistent with cells of human CS7 gastrula (Figure 5D). Human epiblast like cells mainly located in Em-Up, primitive streak cells were including in Em-Mid, while extra-embryonic endoderm cells were separated into Em-Low, YE1 and YE2.

We identified marker genes and regulatory networks that are critical for the molecular annotation of the spatial samples and found there is a considerable conservation between species. For example, the *POU5F1*, *HESX1*, *SOX2* and *NANOG* pluripotency regulatory network were highly activated in embryonic tissues, particularly in the upper layer which are epiblast and its derivatives (Figure 5E), corroborating previous findings (Boroviak et al., 2018) (Nakamura et al., 2016; Niu et al., 2019; Xiang et al., 2019) (Figure S7J and K). Notably, in contrast to that in the rodent model, *NANOG* and *PRDM14* were still highly expressed in the embryonic tissues until the middle and late stages of gastrulation in primate, suggesting a distinct spectrum of pluripotency during gastrulation.

We found that the PS-related genes *TBXT*, *MIXL1* and *FOXA2* showed high expression in the Em-Mid and Em-Low of the epiblast, but was low in the yolk sac endoderm and extra-embryonic mesenchyme (Figure S7J). Similarly, the G4 regulon, consisting of MIXL1, EOMES, FOXA2 and SOX17, were specifically activated in the Em-Low and Em-Mid regions and were enriched with terms such as “Endodermal cell differentiation” and “Regionalization” (Figure 5E). In line with the anatomical annotation, spatial territory of epithelial to mesenchymal transition (EMT) genes demonstrated that epiblast cells at this region undergo an EMT during gastrulation (Figure S7K). Furthermore, the activating activity of the WNT and NODAL signaling pathways were predominantly enriched (Figure 5F and S7K). Em-Low and Em-Mid were distinguished by their relative expression of G1 regulon which were enriched in chromatin remodeling, such as SP1 and KDM4B, suggesting that epigenetics has a pivotal role in lineage commitment (Nicetto et al., 2019; Wang et al., 2018). Therefore, we classified the middle and lower layer of embryo into PS1 and PS2, which may denote gastrulating cells of slightly different status.

Interestingly, the yolk-sac tissues (YE1 and YE2) showed the least similarity to the other dissected regions, as revealed both by UMAP and hierarchical clustering (Figure 5B and S7H). α-fetoprotein (*AFP*), transthyretin (*TTR*) and albumin (*ALB*) were highly expressed in YE1 and YE2, and they also formed co-expression module (Figure S7I and J). The expression of *AFP*, *TTR* and *ALB* was usually regarded as the earliest marker of hepatoblasts (Gualdi et al., 1996; Sheaffer and Kaestner, 2012), while the yolk sac may serves as a pivotal organ for diverse functions of intestine, liver, bone marrow and thyroid even before the matured organs have not been generated at the early embryo development stages (Ross and Boroviak, 2020; Wong and Uni, 2021). Additionally, the apolipoprotein gene family members were also specifically expressed in yolk-sac endoderm, such as *APOA1* (Figure S7J), *APOA2/4* and *APOC2/3* (from DEGs, data not show), which is consistent with the function of the yolk sac as the primary site of apolipoprotein synthesis during early primate and mouse development (Baardman et al., 2013; Cindrova-Davies et al., 2017). Furthermore, as previously reported that primate yolk sac delivers nutrients and oxygen to the embryo early in development (Cindrova-Davies et al., 2017; Dong and Yang, 2018; Exalto, 1995), pathway activity analysis and weighted gene co-expression network analysis (WGCNA) showed that reactome of oxygen release and AMI pathway involving collagen (*COL4A1/2/3/4/5/6*), fibrinogen (*FGA, FGB* and *FGG*) and serpin members (*SERPINC1*) were enriched exclusively in the yolk sac tissues (Figure S7I and K). Notably, we identified FOXA1, FOXA3 and NR1H4 as the key transcript factors in maintaining the identities of yolk sac endoderm (Figure 5E and H) and have the most connections within the regulon group and the node-regulons. Previously, FOXA1 was defined as the visceral/yolk-sac endoderm (VE/YE) marker (Nakamura et al., 2016), hence our analysis further indicated that FOXA1 and FOXA3 correlate with yolk sac endoderm fate specification, whereas FOXA2 were functionally critical for mesoderm and definitive endoderm development (Figure 5E and H).

Cellular fate specification depends on temporally and spatially precise cell communication, so we performed intercellular communication analysis based on ligands, receptors and their cofactors by CellChat (Jin et al., 2021). Although not at single-cell resolution, the cell-cell communication network identified PS1 cells as the dominant communication “hub” (Figure 5G). PS1 cells were mainly enriched with WNT and NOTCH signaling pathway in a paracrine and autocrine manner (PS1 and PS2), which is consistent with their known roles (Figure 5G) in gastrulation (Nakamura et al., 2016; Souilhol et al., 2015). In agreement with previously work in mouse (Ciruna and Rossant, 2001; Sun et al., 1999), we identified that PS1 was the primary FGF source in primate, signaling both in autocrine and paracrine manner with FGF15 − FGFR1, FGF17 − FGFR3 and FGF17 − FGFR1 ligand-receptor pairs to drive the migration of delaminated cells and the differentiation of the mesendoderm.

To uncovered the key sequential signaling events along the process of primate gastrulation, we combined the communication pattern analysis with the previously studied cell events in mouse and human cell models. It’s known that the BMP signaling in the extraembryonic ectoderm in mouse was the source of signaling molecules that transduce the fate of proximal epiblast cells, which formed the primitive streak subsequently (Ben-Haim et al., 2006; Brennan et al., 2001; Chhabra et al., 2019; Rivera-Perez and Hadjantonakis, 2015; Rivera-Perez and Magnuson, 2005). In contrast to mouse, primate embryos form the extraembryonic mesoderm prior to gastrulation, and the BMP signaling was highly enriched in the extraembryonic mesoderm (Figure S7K), suggesting that the extraembryonic mesoderm may serve as a source of signaling molecules in initiation of primate gastrulation. Notably, Hippo-Yap signaling was highly activated in the extraembryonic mesoderm (Figure 5F). Previous work has highlighted the Hippo signaling in regulating early endoderm development during mouse gastrulation (Peng et al., 2019) and the crosstalk between TAZ and SMAD (Varelas et al., 2008), therefore the co-localization of BMP and Hippo/Yap signaling in the extraembryonic mesoderm may hint that Hippo-Yap interacts with BMP signaling to initiate gastrulation of primate. Remarkably, insulin-like growth factor (IGF) signaling was prominently secreted from EXMC2 cells to PS1 cells, and autocrine and paracrine loops of prokineticins (PROK) were established between PS1 and PS2 in monkey gastrulating embryo (Figure 5G and S7L). IGF signaling pathway has been highly conserved and plays an important role in early mesoderm formation of zebrafish and rabbits, which influences the expression of *Otx2*, *FGF8*, *BMP2b*, *Wnt3a* and *Brachyury* (Eivers et al., 2004; Thieme et al., 2012). In addition, IGF1 induced EMT process through the activation of NFκB by the PI3K pathway, then induce SNAIL1 stabilization (Lamouille et al., 2014; Singh et al., 2018). Together, we predicted that BMP, Hippo-Yap and IGF signaling pathway would be imperative to initiate gastrulation in primate, and the communication pattern of CXCL, BTLA, LIGHT and APJ is also from the EXMC to the PS1 specifically (Figure 5G).

### Cellular and molecular dynamic of macaque gastrulation

To further confirm the observed characteristics of germ layer differentiation, the expression of germ layer-specific molecular markers at RNA and protein levels were investigated (Figure 6A and B). Consistent with previous reports, OCT4 (*POU5F1*) was exclusively expressed in the pluripotent cells, especially in the epiblast and ectoderm of early or late gastrulation. GATA6 was specifically detected in the hypoblast by immunohistochemical staining. Notably, we found that the EMT cells were weakly positive for OCT4 and strongly express T, a marker for primitive streak or nascent mesoderm. Interestingly, some mesenchymal cells with the weak expression of T intercalated in the hypoblast (Figure 6 B, white arrowheads), as shown in the histological images analyzed previously (Figure 2 and 3), suggesting a prerequisite for downregulation of the primitive streak genes in forming the definitive endoderm. Furthermore, spatial gene expression showed that *NANOG* still marks pluripotent cells in the epiblast (Figure 6 A and S7J). Taken together our three-dimensional gastrulating model (Figure 6C) provides new insights for the molecular architecture of primate gastrulation.

**Figure 6.**
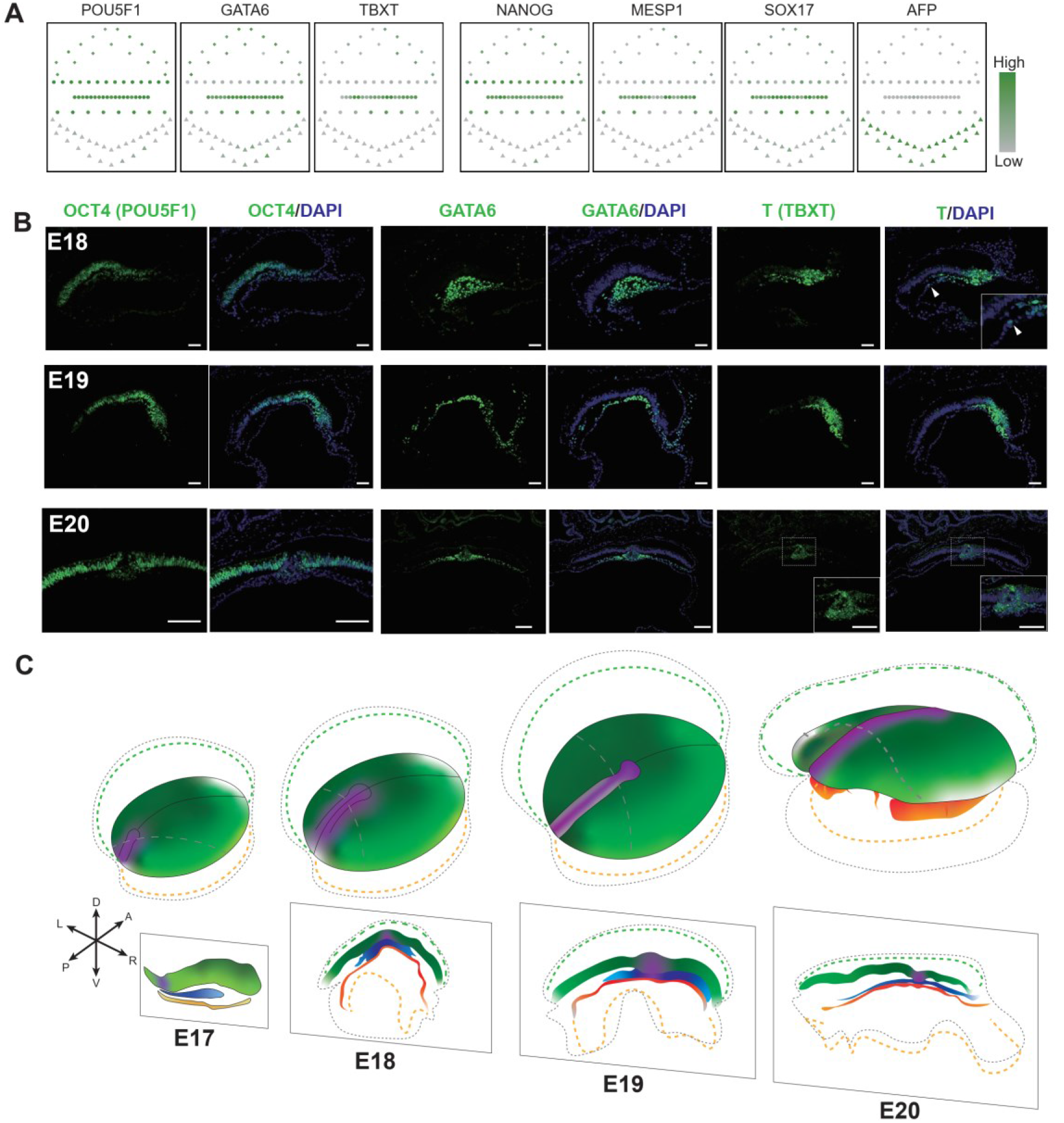
Spatiotemproal cellular and molecular characteristics. A, STGE-plot showing the relative expression of marker genes related to the germ layers and primitive streak in primate gastrulation. Color code represents the relative expressed levels. B, Representative immunofluorescent images of embryos at E18, E19 and E20 for OCT4 (green), GATA6 (green), TBXT (green) and DAPI (blue). White closed arrowheads indicate gastrulating cells that intercalate in the hypoblast. High-magnification images are indicative of the boxed region. Scale bar, 100μm. C, Scheme of monkey gastrulation development. A, anterior; P, posterior; L, left; R, right; D, dorsal; V, ventral. The grey dash lines on epiblast indicate the level of the transversal sections.

### Cross-species spatiotemporal transcriptome analysis of mammalian gastrulation development

Gastrulation is a fundamental morphogenetic process in development, and the key signaling pathways that regulated the establishment of germ layers were evolutionarily conserved, such as the role of WNT, NODAL and BMP signaling pathways in non-human primate and rodent mammalians. However, there are significant difference between primates and mice in both morphology and development rate of gastrulation. To systematically reveal the molecular signatures associated with different mammalian species, we performed cross-species comparative analysis on the spatiotemporal transcriptome in mouse, Cynomolgus monkeys and human gastruloid (STAR Methods). We showed that the states of Cynomolgus monkey at E18 and E20 were resembling to the middle-late stage of mouse gastrulation (Figure 7A and S8A). We aligned the regulons along the developmental time and obtained 5 clusters that correspond to the dynamic patterns of cell fate commitment and regionalization (Figure S8B). All these dynamic regulons (n=451) across gastrulation were subjected to dimensionality reduction analysis. The PCA plot showed that the cell types between monkey and mouse were clustered together, suggesting a germ lay consistency (Figure 7B). The PS1 and PS2 in macaque gastrulating embryos were close to the mouse mesoderm and endoderm, respectively. To explore the species-specific molecular mechanism underlying the germ layer segregation, we defined a species/spatial domain-specificity score (SSS) based on Jensen-Shannon divergence (STAR Methods). The activities of most regulons (more than 50%) were similar during germ layer formation in mice and monkeys, suggesting that the major gastrulation events are conserved and common molecular regulatory mechanisms are shared between species (Figure 7C-E, S8C-E, Table S3). Next, we looked into the species difference in germ layers and whole embryonic tissues (Figure 7C-F). For quantitative visualization of species specificity, relative SSS of regulons were plotted in ternary plot (Figure 7F). The activity of KLF7 were preferred in mouse, especially in the ectoderm and the endoderm differentiation (Figure 7C, E and F), whereas the ELF3 were enriched in the ectoderm of macaque (Figure 7C and F). Given that *Klf7* had inhibitory activity on mesoderm (Gao et al., 2015) and *Elf3* was a negative regulator of epithelial-mesenchymal transition in gastrulation and tumor metastasis (Scheibner et al., 2021; Yeung et al., 2017), this might imply that mice and non-human primates use functionally similar but different regulatory factors in the maintenance of the ectoderm epithelium.

**Figure 7.**
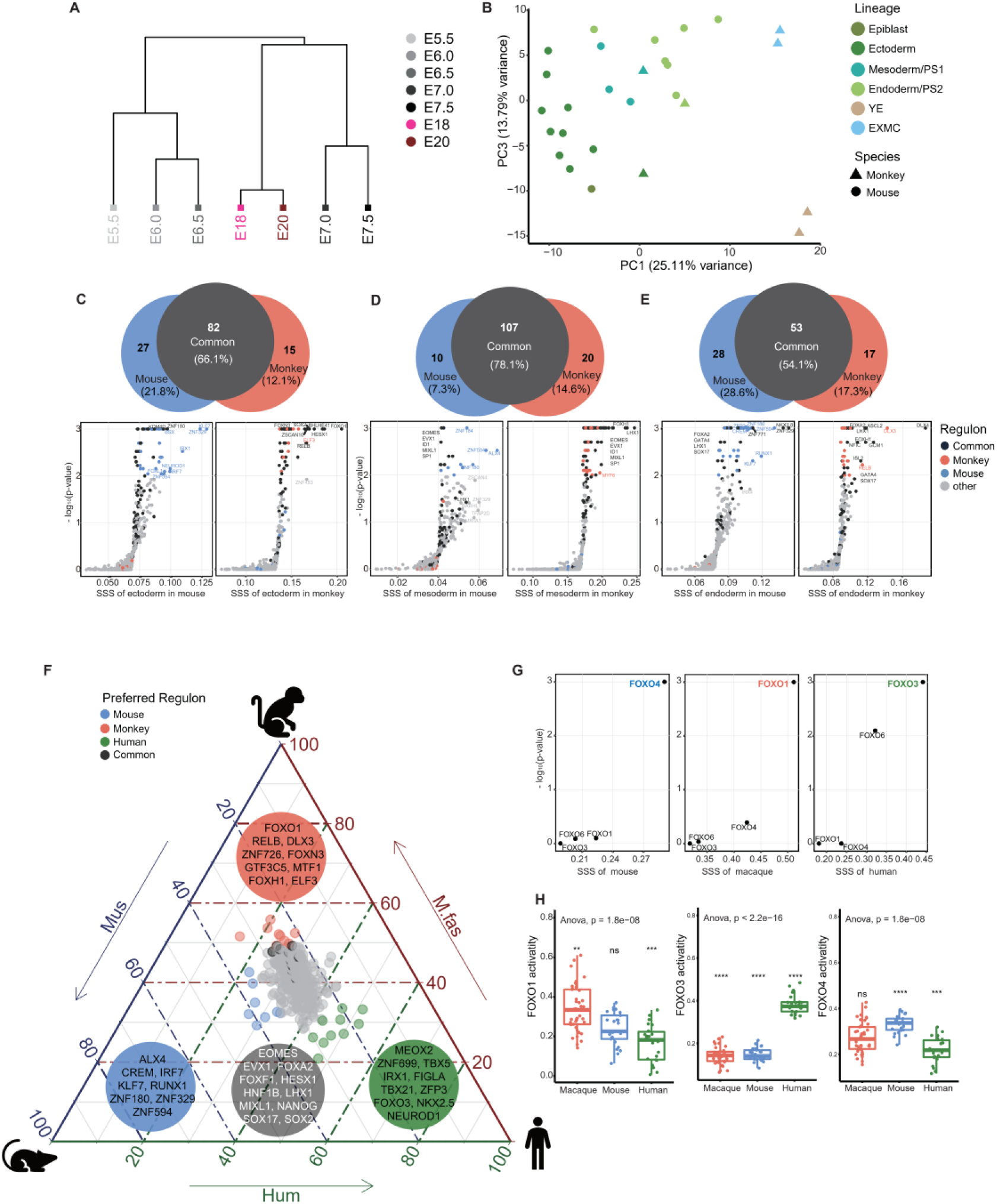
Cross-species spatial transcriptomic analysis reveals gastrulation developmental differences among mice, monkeys and humans. A, Hierarchical clustering analysis of mouse and monkey whole embryos in different gastrulation stages based on averaged dynamic regulons (n=451). B, PCA analysis showing the lineage difference between mouse and monkey. YE, yolk-sac endoderm; EXMC, extra-embryonic mesenchyme tissues. C-E, Species specific regulons between mouse and monkey in ectoderm, mesoderm and endoderm embryogenesis, respectively. Venn plot showed the lineage species specific and common regulons. The scatter plot showed the significant and specific regulons in different species germ layer. SSS, species/spatial domain-specificity score. Color code the specific type of regulons. F, Ternary plot of the most divergent species-specific regulons in gastrulation between mouse, monkey and human. Top 10 regulons were colored and listed based on species - specificity score (SSS). G. Species-specificity score of FOXO regulons between mouse, monkey and human gastrulation embryonic tissue. H, The activities of FOXO regulons between mouse, monkey and human gastrulation embryonic tissue.

Interestingly, we found that Forkhead box transcription factor of subclass O (FOXO) family showed species-dependent activity. Specifically, we showed that FOXO1 was exclusively activated in macaque gastrulating embryos and mouse embryos exhibited the highest levels of FOXO4 regulons, while FOXO3 activity was substantially elevated in human (Figure 7G-H). In line with that FOXO1 regulates core pluripotency factors in human and mouse embryonic stem cells by promoting the expression of *OCT4* and *SOX2* (Zhang et al., 2011), we found that FOXO1 have a strong positive correlation with stem cell pluripotency genes *SOX2* and *ZSCAN10*, which enriched in negative regulation of cell differentiation and transcriptional regulation of pluripotent stem cells (Figure S8F). Furthermore, we identified the targets of FOXO1/3/4 and their top related genes. They also showed species differences with FOXOs expression, including *TERT* (telomerase reverse transcriptase), *BCL6* and *TEF* (thyrotroph embryonic factor) in mouse, macaque and human embryos respectively (Figure S8H-J) (Chakravarti et al., 2021; Tang et al., 2002; Yamashita et al., 2014; Yang et al., 2019). Together, our data indicate that FOXOs may play a vital role in interspecies differences during gastrulation. Considering the prominent role of FOXO family in metabolic homeostasis and organismal longevity, the species-specific mechanisms may imply profound effect of FOXO in even beyond embryo gastrulation.

## Discussion

In this study, we have provided an anatomical atlas and molecular database spanning the entire gastrulation to the early organogenesis period of Cynomolgus monkey development. This analysis was a critical step in understanding of human embryology within similar time-window, which remain largely unknown due to technical difficulties and ethical issues. First, we have captured a series of high-resolution tissue anatomical sections from early gastrulation (E17) to early organogenesis (E21), revealed the dynamic changes of cell histomorphology and cell proliferation kinetics, and discovered the critical role of cell migration and EMT/MET transformation in the process of germ layers formation. Secondly, the three-dimensional model of primate embryos was reconstructed to analyze the dynamics of gastrulation from a holistic perspective. Furthermore, the spatial transcriptome of macaque gastrula has been performed. We identified the key regulons and their interaction networks in different cell types in the process of germ layer segregation. Finally, we have explored cross-species differences in mouse, Cynomolgus monkeys and human by comparatively analyzed the spatiotemporal transcriptomes.

Gastrulation is a period of rapid growth, proliferation and differentiation. Several morphological and genetic reports in mouse have shown that the cell cycle progression is coordinated with transcription, cell migration, and cell differentiation (Abe et al., 2013; Mitiku and Baker, 2007; Snow, 1977; Tam and Behringer, 1997), while little is known about this in primate gastrulation. We have applied detailed histological analyses to assess the growth rate by determining the cell number increase, and highlighted the cell proliferation kinetics in primate gastrulation. However, the tissue- and embryo-wide division patterns in primate gastrulating embryos needs further to be studied through genetic markers labeling and spatial omics, because the rate of proliferation differs significantly across the different regions of the epiblast in mouse at gastrulation stages (McDole et al., 2018; Snow, 1977).

Although the morphology of primate embryonic development has been studied for at least a century (de Bakker et al., 2016; Grobstein, 1985; Hendrickx, 1972; Luckett, 1978), detailed anatomical imaging with high temporal resolution in primate gastrulating embryos are lacking. We provided an atlas with sequential stages and serial sections of gastrulating embryos in 2D and 3D, which was fundamental to reveal cellular morphological changes during ectoderm, mesoderm and endoderm formation. The results of this study showed that mesenchymal like cells were found intercalated in hypoblast individually, not as a coordinated sheet to displace the original hypoblast (Figure 2 and 3). And immunohistochemical staining indicated that embedded mesenchymal cells weakly expressed the primitive streak marker T (Figure 6B), implying that the original hypoblastic cells could become integrated into the newly formed embryonic endodermal layer and EMT/MET play important roles in the embryogenesis of primate endoderm.

Remarkably, from the spatial transcriptome of monkey gastrulating embryos, FOXA subfamily was found to have important biological functions in embryonic and extraembryonic endoderm specification (Figure 5E and H). From the evolution and phylogeny view, FOXA genes have a conserved role in the development of the derivatives of the primitive gut (Hannenhalli and Kaestner, 2009; Lai et al., 1991), whereas the functional diversity among FOXA proteins with the spatio-temporal expression patterns were not clear. In mouse, neither *Foxa1* nor *Foxa3* inactivated mutants exhibit any early phenotype (Grapin-Botton and Constam, 2007; Kaestner et al., 1998), whereas they can compensate for the loss of *Foxa2* in the null mutants, which allows hindgut, but not foregut and midgut formation (Dufort et al., 1998; Monaghan et al., 1993; Sasaki and Hogan, 1993). In detail, the *FOXA1* was not just as the yolk-sac endoderm marker (Nakamura et al., 2016), but also regulate yolk sac endoderm fate specification with *FOXA3*. On the other hand, *FOXA2* had a pivotal role in cell fate commitment of embryonic endoderm, which was consistent with the finding in single cell transcriptome of human (Tyser et al., 2020). Indeed, the molecular map of cell populations at defined positions varies. Our Geo-seq approach may not be able to separate the mesoderm and endoderm cells at single-cell resolution, therefore results in a relatively similarity at transcription level to be defined as PS1 and PS2, respectively (Figure 5E). Further research should be undertaken to explore the spatiotemporal molecular architecture at single cell resolution and to investigate the mechanism of gut endoderm formation in nonhuman primates.

With the spatial transcriptome data of Cynomolgus embryos, we can now glean the primate-specific molecular program by cross-species comparative analysis. We defined species-specific regulons. As expected, most regulons were shared in mice and monkey embryos during germ layer formation, suggesting that the gastrulation event were evolutionally conserved. Interestingly, we found FOXO family showed species-specific usage during gastrulation. FOXO1, FOXO3 and FOXO4 were enriched in macaque, human and mouse respectively (Figure 7G-H), demonstrating that FOXOs may have crucial functions in interspecies evolution. As a key transcription factor to integrate different signals from the insulin/ insulin-like growth factor 1 (IGF-1) signaling pathway, target of rapamycin (TOR) signaling, AMP-activated protein kinase (AMPK) pathway and Jun N-terminal kinase (JNK) pathway (Fontana et al., 2010; Lin et al., 1997; Ma and Gladyshev, 2017; Sun et al., 2017; Tian et al., 2017) to participate in a wide range of important cellular processes such as cell cycle arrest, apoptosis, and metabolism besides its function in stress resistance and longevity (Calissi et al., 2021; Eijkelenboom and Burgering, 2013; Golson and Kaestner, 2016). The preferred cellular mechanisms of FOXO family during early embryo development and beyond is awaiting further investigation.

In summary, our study provides a morphological and molecular atlas for illustrating the dynamics of key processes and regulatory mechanisms during three germ layers formation in primate embryos. Knowledge gained from this work should be valuable for evaluating the interspecies difference and setting a reference for in vitro mimic of monkey embryogenesis. Our work also provide a reasonable working model for human gastrulation. To fully dissect the primate gastrulation, additional studies will be needed to build the single-cell spatiotemporal molecular maps with implementing genetic manipulation and lineage tracing on monkey models.

## SUPPLEMENTAL INFORMATION

Supplemental Information includes Supplemental Experimental Procedures, eight figures, and three tables and can be found with this article online at ..

## ACKNOWLEDGMENTS

We thank W. Ji for critical discussions, S.Suo for help of data analysis, and J. Xu, J.Zhang, M.Wang, L.Qin for experimental support. This work was supported in part by National Key R&D Program of China (2018YFA0801402, 2018YFA0107200, 2019YFA0801400), the “Strategic Priority Research Program” of the Chinese Academy of Sciences (XDA16020404 and XDA16010308), National Natural Science Foundation of China (31871456, 32100483), Guangdong Basic and Applied Basic Research Foundation (2019B151502054, 2019A1515110985).

## AUTHOR CONTRIBUTIONS

N.J. and G.P. conceived the study. G.P., N.J. and W.S. supervised the project. G.P., N.J., W.S. and G.C. designed the experiments. G.C. and S.F. conducted the histology and IF and analyzed the data, G.C. and L.W. contributed to the 3D reconstruction of the histological sections, W.S., Y.Y., X.H., X.L., Y.D. and P.Z. performed samples collection, G.C. and G.P. performed the Geo-seq experiments and analyzed the spatial transcriptome data, K.T. and J.C. performed tissue staining, G.C., G.P., P.P.L.T. and N.J. wrote the paper with the help of all other authors.

## STAR Methods

### KEY RESOURCES TABLE

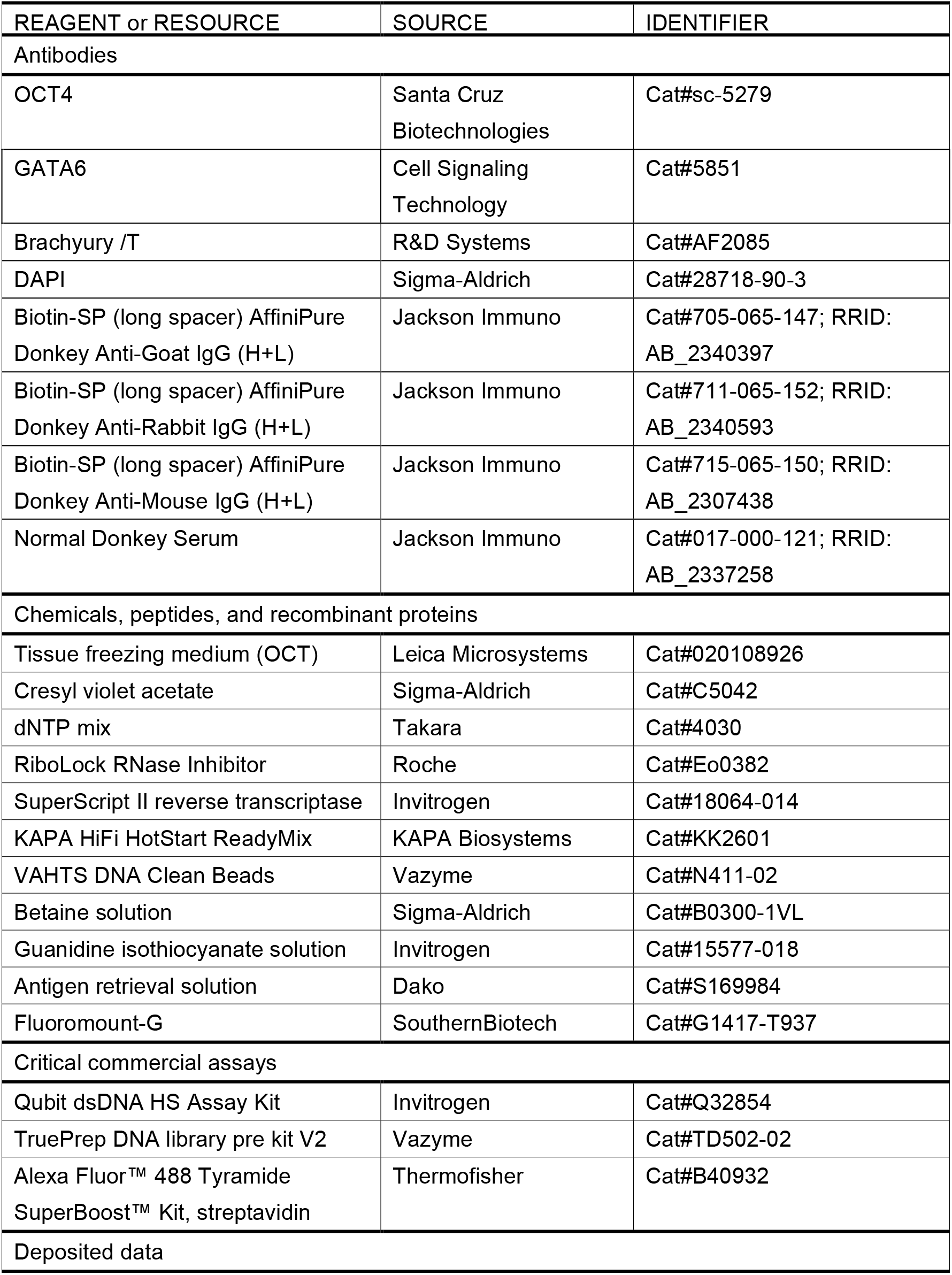
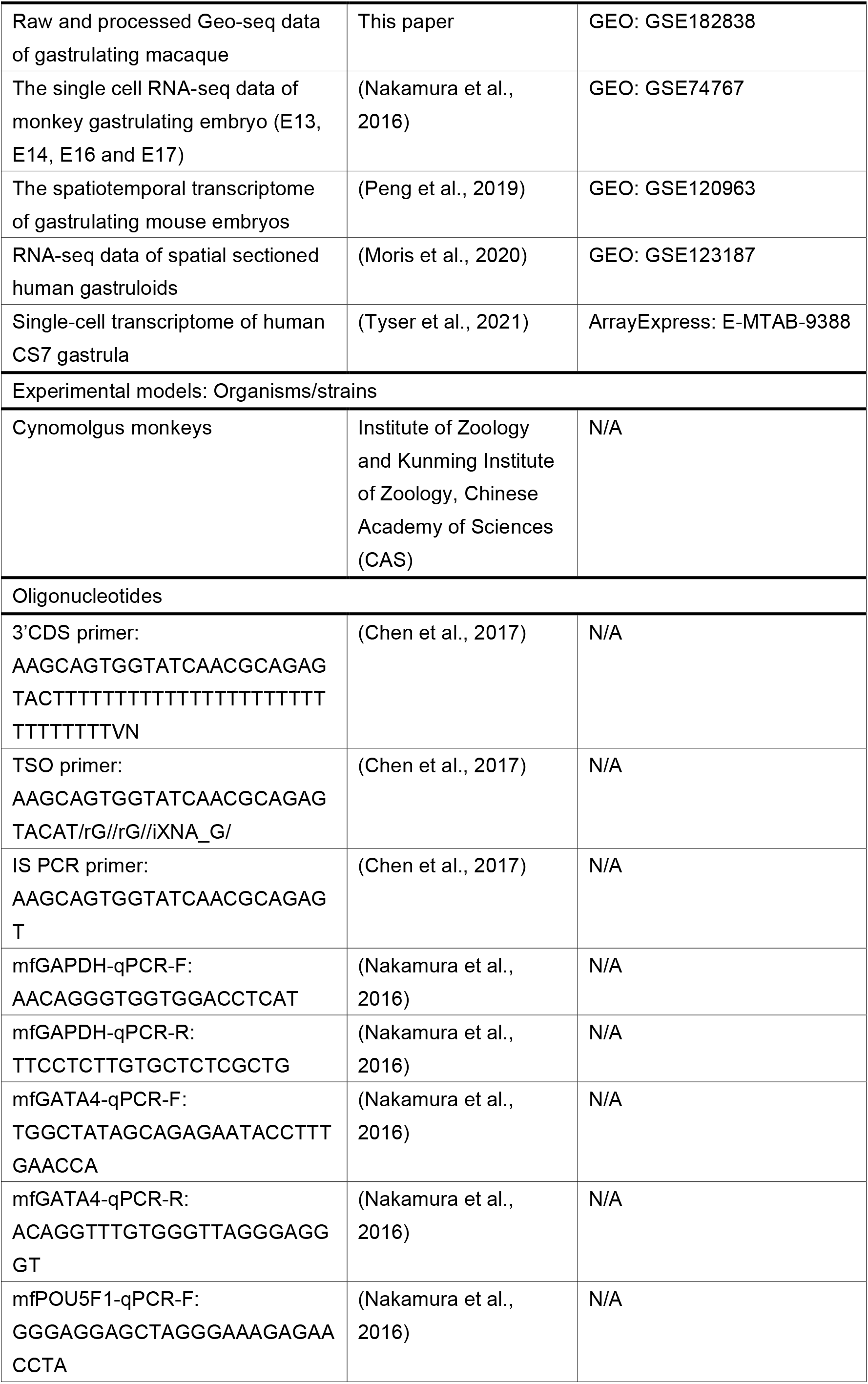
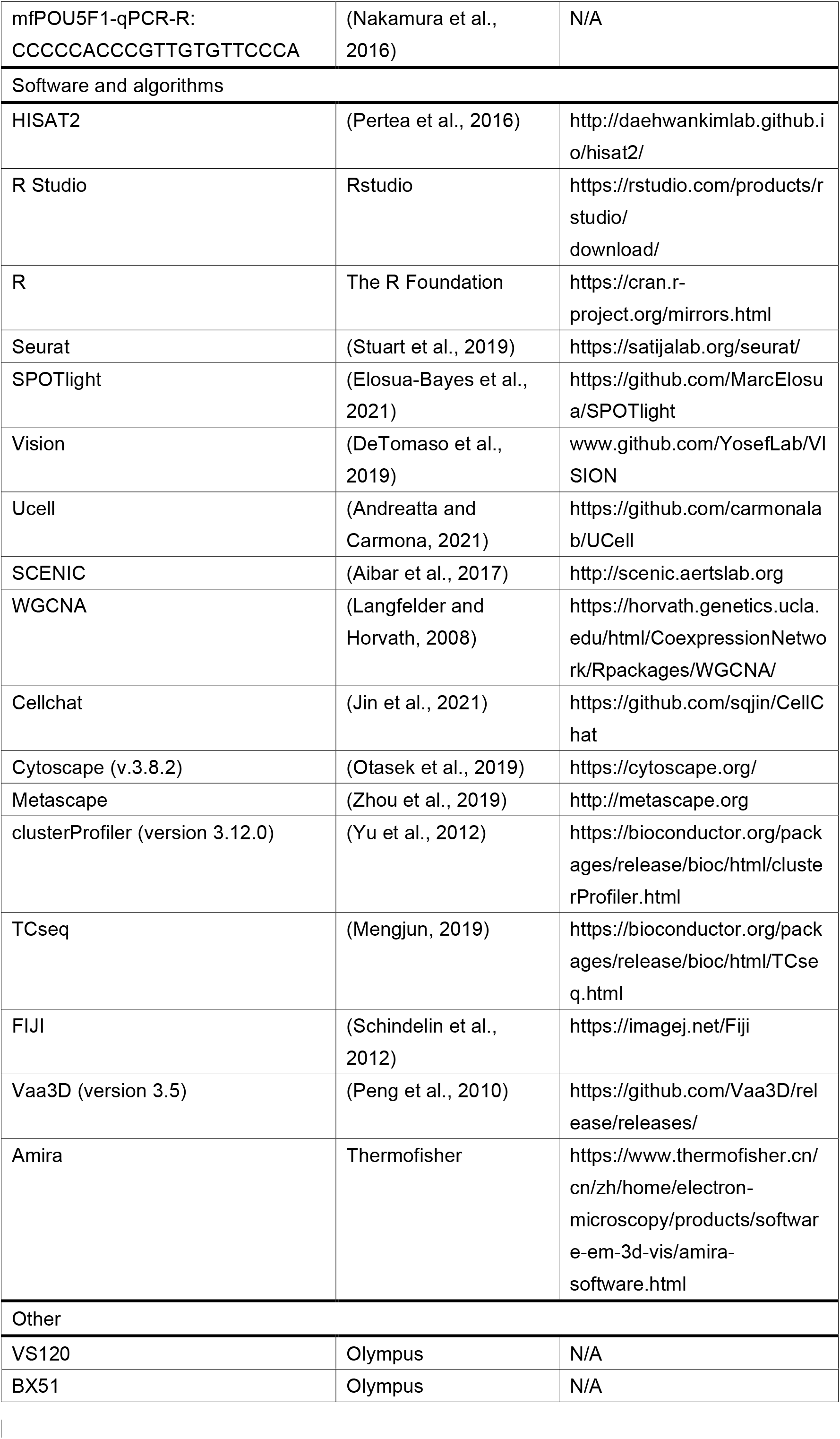

### RESOURCE AVAILABILITY

#### Lead contact

Further information and requests for resources and reagents should be directed to and will be fulfilled by the Lead Contact, Guangdun Peng.

#### Materials availability

Unique materials generated in this study are available from the Lead Contact without restriction.

#### Data and code availability

The Cynomolgus monkey spatiotemporal RNA-seq raw and processed data have been deposited in the Gene Expression Omnibus database under accession number GSE182838, and also can be downloaded from our website. Custom code and scripts are available from methods details and github. All other data supporting the findings of this study are available from the corresponding author upon request.

### EXPERIMENTAL MODEL AND SUBJECT DETAILS

All monkey experiment procedures were under the guidance of the Ethics Committee of the Institute of Zoology and Kunming Institute of Zoology, Chinese Academy of Sciences (CAS). The gastrulating embryos of Cynomolgus monkeys (*Macaca fascicularis*) from natural conception and in vitro fertilization (IVF) were both used in this study. The procedures in Cynomolgus monkeys for oocyte collection, intra-cytoplasmic sperm injection, pre-implantation embryo culture, and transfer of pre-implantation embryos into foster mothers were performed as described previously (Niu et al., 2019; Yamasaki et al., 2011). The day when the intra-cytoplasmic sperm injection was performed was designated as embryonic day (E) 0. For the detection of pregnancy of early post-implantation embryos, implanted embryos were monitored by ultrasound scanning around E14 and the implanted uterus was surgically removed and bisected for the isolation of embryos.

For naturally mated embryos, health adult female and male monkeys aged from 7-8 years were individually kept in an animal room with humidity at 40-70% and temperature at 18-26 °C and exposed to a 12:12 light–dark cycle. The female monkeys were observed at least two normal menstrual cycles confirmed by virginal bleed before being allowed to mating with male monkeys. The first day of bleeding was defined as day 1 of menses once menstrual cycle start. Then venous blood was collected at 09:00 AM on menses days 7 to 18, and a male monkey was allowed to mating with the female monkey. Blood serum was separated by centrifugation and concentrations of E2 and P4 were assayed using a chemiluminescent immunoassay. Once the peak of E2 was detected, then the next day was defined as the day of ovulation and fertilization, and the concentration of blood E2 and P4 were continuously measured. The pregnancy was confirmed by the low level of E2, increased level of P4 and ultrasonography.

### METHOD DETAILS

#### Monkey embryo H&E staining and immunofluorescent analysis

Monkey post-implantation embryos by IVF procedure were harvested at different stage from embryonic day 17(E17) to E21 every day (at least two embryos in each stage), and whole embryos were fixed in 4% paraformaldehyde (PFA) at 4°C overnight. The procedures of paraffin embedding were carried out, and the embedding embryos were transversely sectioned at a thickness of 5 μm. The sections were stained by H&E staining and immunofluorescent staining as previously reported (Feng et al., 2017). In brief, after dewaxed and rehydrated, the slides were stained in hematoxylin solution for 30 sec and counterstained in eosin solution for 10 sec. For immunofluorescence staining, the sections were incubated with primary antibody in hybridization buffer overnight at 4 °C after blocked, and incubated with secondary antibody for 1 hour (h) at room temperature. The sections were counterstained with 4′,6′-diamidino-2-phenylindole (DAPI) and mounted with mounting medium. Bright field images and fluorescent images were collected on an Olympus VS120 and Olympus BX51 microscope respectively, and aligned manually using Adobe Photoshop CC.

#### Determination of gastrulation cell number and mitotic activity

The number of cells contained in epiblast/ectoderm, mesoderm and hypoblast/endoderm were determined by counting nuclei on every section of the gastrulating embryos. The total score was then adjusted by applying Abercrombie’s correction formula (Abercrombie, 1946) to give an estimate of the actual cell number. In detail, the average number of cells per section (*P*):

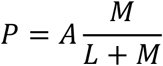

*A* is the crude count of number of nuclei fragments, *M* is the thickness of the section (*M*=5μm in this study), and L the average length of the nuclei (*L*=10μm in this work).

Cell cycle times can be calculated from the equation (Snow, 1977), *C_t_* means the cell number at *t* stage:

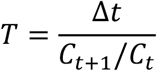

#### 3D reconstructions of gastrulating monkey embryos

The Vaa3D (version 3.5) and Amira (version 6.0.1) were used for the 3D reconstruction process, consisting of alignment, segmentation and visualization (de Bakker et al., 2016; Peng et al., 2010). In brief, grey scale processing was performed by Photoshop CC (Adobe Systems), and alignment of the stitched serial images was mainly implemented by manual after automatically adjustments. The aligned 3D images were segmented into epiblast/ectoderm, primitive streak, hypoblast/endoderm and mesoderm. Then, triangulated surface files were made using the *SurfaceGen* function, and triangle reduction (*Simplify*) and surface smoothing (*SmoothSurface*) were also applied for visualization. Besides, the Vaa3D was also applied to aligned sections to visualize the 3D structure of specific tissues.

#### Spatial molecular profiling of monkey embryos

The spatial transcriptome of gastrulating embryos was obtained according to Geo-seq protocol with minor modifications (Peng et al., 2019). Briefly, whole embryos (two monkey embryos at E18 and one at E20 from natural conception) were embedded in OCT (Leica Microsystems, catalogue no. 020108926), and cryo-sectioned in the coronal plane serially from the anterior region to the posterior region at a thickness of 20 μm. Sections were transferred onto LCM PEN membrane slides, fixed immediately by ethanol and stained with 1% cresyl violet acetate in 75% ethanol solution (Sigma-Aldrich). The embryonic tissue was divided into upper, middle and lower layer, and the extraembryonic tissue was divided into extra-embryonic mesenchyme 1/2 and yolk-sac endoderm 1/2 based on the good morphology sections. Populations of 5-50 cells were collected by laser microdissection, and total RNA pellets were dissolved in lysis solution, followed by reverse transcription using SuperScript II reverse transcriptase (Invitrogen), whole transcription amplification with KAPA HiFi HotStart ReadyMix (2X; KAPA Biosystems). The PCR product of LCM samples were used for automated single-cell RNA-Seq library construction based on the Bravo robot station(Cui et al., 2019). In brief, PCR product were purified using 0.75X AMPure XP beads (Agencourt), quantified with Qubit dsDNA HS Assay Kit (Thermo Fisher) on Envision (PerkinElmer), and cDNA library was constructed by DNA Library Prep Kit V2 for Illumina (Vazyme) and sequenced on an Illumina Nova 6000 instrument using a 150 bp paired-end-reads setting.

### QUANTIFICATION AND STATISTICAL ANALYSIS

#### Processing of RNA-seq data

Sequencing quality of raw sequencing data was evaluated by fastp (Chen et al., 2018). The genome sequence Macaca_fascicularis_5.0 for Cynomolgus monkeys (*Macaca fascicularis*) were obtained from the Ensemble (https://asia.ensembl.org/index.html). Reads were mapped to the Macaca_fascicularis_5.0 genome assemblies by HISAT2 (Pertea et al., 2016) using default settings. Mapping ratio was calculated based on the number of mapped reads and total reads for each sample. All mapped reads were processed by StringTie(Pertea et al., 2016) to quantify gene expression levels (measured by TPM and FPKM, Transcripts Per Kilobase per Million mapped reads and Fragment Per Kilobase per Million mapped reads respectively) using default parameters. Seurat package (version 3.1.5) (Stuart et al., 2019) was applied to dimensionality reduction analysis in R (version 4.0.3). Gene expressed in at least two samples across all samples were selected for further analysis. After filtering, data in each sample were normalized, the 5,000 most variable genes were identified with *vst* method, and the expression levels of these genes were scaled before performing Principal Component Analysis (PCA) in variable gene space. Next, 5 principal components were used for graph-based clustering (resolution = 2) and UMAP dimensionality reduction was computed using *RunUMAP* function with default parameters. The Spearman correlation coefficient (SCC) of spatial domain between embryonic replicates or developmental stages were calculated based on average expression of all gene.

#### Identification of differentially expressed genes (DEGs) and clustering analysis

Gene expression levels for each sample were log_2_ (TPM+1) transformed and then combined 5,000 most variable genes with top 150 highest and lowest Principal Component (PC) loading genes (by using *FactorMineR* in R) from PC1-5 to identify the DEGs. In total, 1,305 genes were identified as DEGs. Finally, unsupervised hierarchical clustering was performed using the *hclust* function Ward’s method (ward.D2) to determine the final spatial domains of embryonic and extraembryonic tissues based on the expression profile of DEGs.

#### Integrating scRNA-seq and spatial transcriptome

To investigate the architecture of the cell-type distribution in spatial transcriptome, we integrate both single-cell and spatial transcriptomics data of monkey gastrulating embryo *in vivo* to infer the spatial locations of different cell types in a tissue. The single cell RNA-seq data of monkey gastrulating embryo (E13, E14, E16 and E17) from GSE74767(Nakamura et al., 2016), and apply *FindAllMarkers* function of Seurat pipeline to identify cluster markers. SPOTlight, a deconvolution algorithm that built upon a non-negative matrix factorization (NMF) regression algorithm was used for cell-type deconvolution in spatial transcriptome (Elosua-Bayes et al., 2021). In brief, SPOTlight factorizes the normalized scRNA-seq gene expression matrix into two lower dimensionality matrices using NMF, and cell-level topic distribution matrix is used to learn the cell-type specific topic profiles through a Non-Negative Least Squares regression (NNLS). The weights of each cell-type specific topic profile represent the cell-type proportions across all samples in spatial transcriptome. Furthermore, we used the averages of single-cell detectable gene expression in the same annotated cell type and spatial samples at the same location to calculate the SCC.

#### Pathway activity analysis and epithelial-mesenchymal transition scores analysis

To identify the key biological properties of spatial samples, Vision (DeTomaso et al., 2019; Zhang et al., 2020) and Ucell (Andreatta and Carmona, 2021) were applied to perform functional enrichment analyses. The homologous target genes of WNT, BMP, FGF, NODAL and Hippo/Yap signaling pathways between monkey and mouse gastrulating (Peng et al., 2019) were used as signatures of *vision* analysis. In addition, we input the gene signatures of MSigDB, which curated by the Broad institute, to explore the transcriptional effects of regions. To evaluate the epithelial–mesenchymal transition (EMT) of epiblast cells, scoring EMT based on the Mann-Whitney U statistic (Andreatta and Carmona, 2021). The EMT-inducing transcriptions factors (EMT-TFs) zinc-finger E-box-binding family (ZEB1 and ZEB2), basic helix–loop–helix transcription factors (TWIST1 and TWIST2) and snail family transcriptional repressor (SNAI1 and SNAI2) were included as signature of mesenchymal state. The markers associated with the mesenchymal state vimentin (VIM), fibronectin and β1 (FN1) and matrix metalloproteinase (MMP2) and the markers related to the epithelial state E-cadherin (CDH1), grainyhead like transcription factor (GRHL2), epithelial cell adhesion molecule (EPCAM), Occludins (OCLN), Claudins (CLDN4 and CLDN7) and cytokeratins (KRT19) were also add into Ucell analysis (Chakraborty et al., 2020; Dongre and Weinberg, 2019; Lamouille et al., 2014). The EMT score is the difference of epithelial score minus mesenchymal score (E-M), and the higher EMT scores, the more epidermal-like state.

#### Reconstruction of regulons in embryonic and extraembryonic tissues

For inferring Cynomolgus monkey’s gene regulatory network (GRN) using SCENIC (Aibar et al., 2017), common genes listed in the Cynomolgus monkeys–humans one-to-one annotation table were used(Nakamura et al., 2016). 17,019 common genes were first filtered to exclude all genes detected in fewer than two samples, and then were used to identified TF-gene co-expression modules. Subsequently, those modules are refined via RcisTarget by keeping only those genes than contain the respective transcription factor binding motif. Finally, SCENIC found 323 regulons in monkey gastrulating embryos, whose regulatory activities were represented by AUCell values. To identify regulons associated with spatial domains of gastrulating monkey embryos, we used the regulon activity score for each sample to perform PCA, heatmap display and network analysis (Peng et al., 2016; Peng et al., 2019). Regulons shared in 250 most variable regulons and top 100 highest and lowest PC loading regulons from PC1-3 were defined as specific regulons. For network visualization, the regulon was assigned to the tissue with highest average regulon activity score. The SCC was calculated in the same lineage, and the absolute value was greater than 0.95 for further analysis. The edge weights were proportional to the SCC values of two correlated nodes.

#### Co-expression gene network and interaction analysis

To identify biologically relevant patterns of spatial samples, we performed weighted gene co-expression network analysis using R package WGCNA (Langfelder and Horvath, 2008). Briefly, a soft power threshold of 10 was set to constructed unsigned network and the minimum module size was set to 10 genes. Eight gene modules with significant correlation in the seven spatial cell populations were used to draw gene modules for each cell type. Only top 30 genes with highest 100 correlation coefficient values in each module were kept when visualizing the modules with Cytoscape (v.3.8.2)(Otasek et al., 2019).

To quantitatively infer and analyze intercellular communication networks, Cellchat was applied to predict potential interaction probability scores (Jin et al., 2021). Human gene names were converted to Cynomolgus monkey genes using orthology data from monkeys–humans one-to-one annotation table, with 17,542 genes in common between the two species (Nakamura et al., 2016). We use the tissue type of spatial sample based on regulon activity clustering as label information, and calculated an aggregated interaction probability by Cellchat based on the expression level of ligand–receptor pairs.

#### Functional enrichment analysis and phenotype analysis

We applied Metascape (Zhou et al., 2019) (http://metascape.org), KOBAS-i (KOBAS intelligent version, version 3.0) (Bu et al., 2021) and R package clusterProfiler (version 3.12.0) (Yu et al., 2012) to perform Gene Ontology and pathways enrichment analysis for each group of DEGs and regulon TFs. The TFs which display strong gastrulation phenotypes from Mouse Genome Information (MGI) database were highlighted and visualized in the regulon network.

#### Cross-species comparative analysis

The spatiotemporal transcriptome of gastrulating mouse embryos was generated in our previous work (Peng et al., 2019) (GSE120963), and used the RNA-seq data of spatial sectioned human gastruloids (Moris et al., 2020) (GSE123187) and single cell transcriptome (E-MTAB-9388) to assess human gastrulating embryos. For comparison of the gene expression among mouse, Cynomolgus monkeys and human, ortholog genes of these three species from Ensemble were used for further analysis (Nakamura et al., 2016). To reveal the temporal pattern of time course sequencing data, the TCseq package (Mengjun, 2019) was performed to classify the dynamic regulons into various types of clusters, with the genes in each cluster were then processed for functional enrichment analysis. Then the total of 451 different regulons across five stages of gastrulation were used for hierarchical clustering and PCA analysis.

To quantify the species germ layer specificity of a regulon, an entropy-based strategy was modified from regulon specificity score (RSS) (Peng et al., 2019; Suo et al., 2018). All 490 regulon gene lists which calculated by SCENIC in monkey gastrulating embryos were reanalyzed in natural logarithms transformed expressed matrix of three species by AUCell package (version 1.16.0). Then the Jensen–Shannon divergence (JSD) algorithm was used to identify species-spatial domain-specific regulons by philentropy package (version 0.5.0) (Drost, 2018). Finally, for distribution of each regulon *P_1_* and predefined pattern *P_2_*, the species/spatial domain-specificity score (*SSS*) between *P_1_* and *P_2_* was defined by converting JSD to a similarity score:

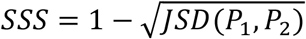

To evaluate the significance of regulon specificity in each species spatial domain, the regulon-activity score was permuted across all samples 1,000 times, then calculated *SSS*, and finally a permutation *p* value was calculated as the number of times that *SSS*_permutation_ > *SSS*_true_ divided by 1,000. Only the significant regulons (*P* < 0.01), which were specifically activated in species-spatial domains, were kept for downstream analysis. Ternary plot was applied to show the proportion of *SSS* mice, monkeys and humans.

## SUPPLEMENTAL INFORMATION

**Figure S1.**
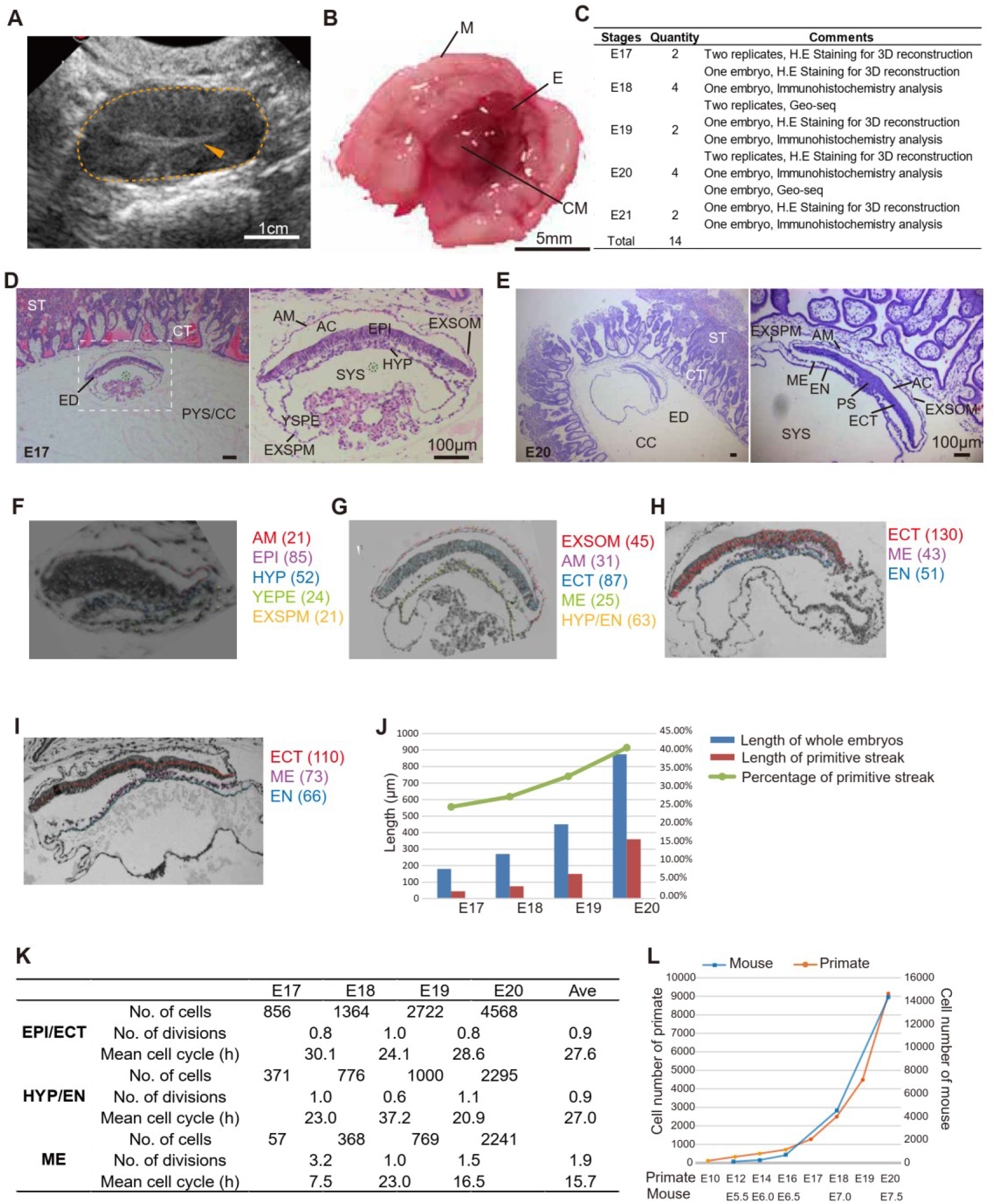
Macaque gastrulating embryos. A, Ultrasound diagnosis of the monkey’s recipient uterus for the implantation of transplanted embryos at E17. Dashed circles indicate the uterus and arrowheads indicate the chorionic cavity. Scale bars, 1 centimeter. B, Images of dissected monkey uterus at gastrulation stage. M, myometrium; E, endometrium; CM, chorionic membranes. Scale bars, 5 millimeters. C, Summary of collected embryos during gastrulation. D, Haematoxylin and eosin staining of the section of replicate embryo at E17. The image at right is a higher magnification of the area boxed on the left. ST, syncytiotrophoblast; CT, cytotrophoblast; PYS, primary yolk sac; CC, chorionic cavity; ED, Embryonic disc; AC, amniotic cavity; AM, Amnion; EPI, epiblast; HYP, hypoblast; YSPE, yolk-sac parietal endoderm; EXSPM, extraembryonic splanchnic mesoderm; EXSOM, extraembryonic somatic mesoderm; SYS, secondary yolk sac. Scale bar, 100μm E, Haematoxylin and eosin staining of the section of replicate embryo at E20. The left is the lower magnification images. Scale bar, 100μm F-I, The number of nuclei fragments contained in epiblast/ectoderm, mesoderm and hypoblast/endoderm were counted on every section of the E17 (F), E18 (G), E19 (H) and E20 (I) embryos. AM, Amnion; EPI, epiblast; ECT, ectoderm; ME, mesoderm; HYP, hypoblast; EN, endoderm; YSPE, yolk-sac parietal endoderm; EXSPM, extraembryonic splanchnic mesoderm; EXSOM, extraembryonic somatic mesoderm. J, The percentage of length of primitive streak. The length of embryo was assessed by total sections, while using sections with gastrulation area to evaluate the length of primitive streak. K, Cell numbers and mitotic activity of germ layers of gastrulating monkey embryos. EPI, epiblast; ECT, ectoderm; ME, mesoderm; HYP, hypoblast; EN, endoderm. L. Cell numbers of gastrulation in mouse and primate embryos. The cell number of primate embryos before E16 were got from in vitro cultured human embryos (Xiang et al., 2019), and the cell number of primate embryos during gastrulation were estimated by histological analysis. The cell number of mouse embryos were adapted from histological determination (Snow, 1977).

**Figure S2.**
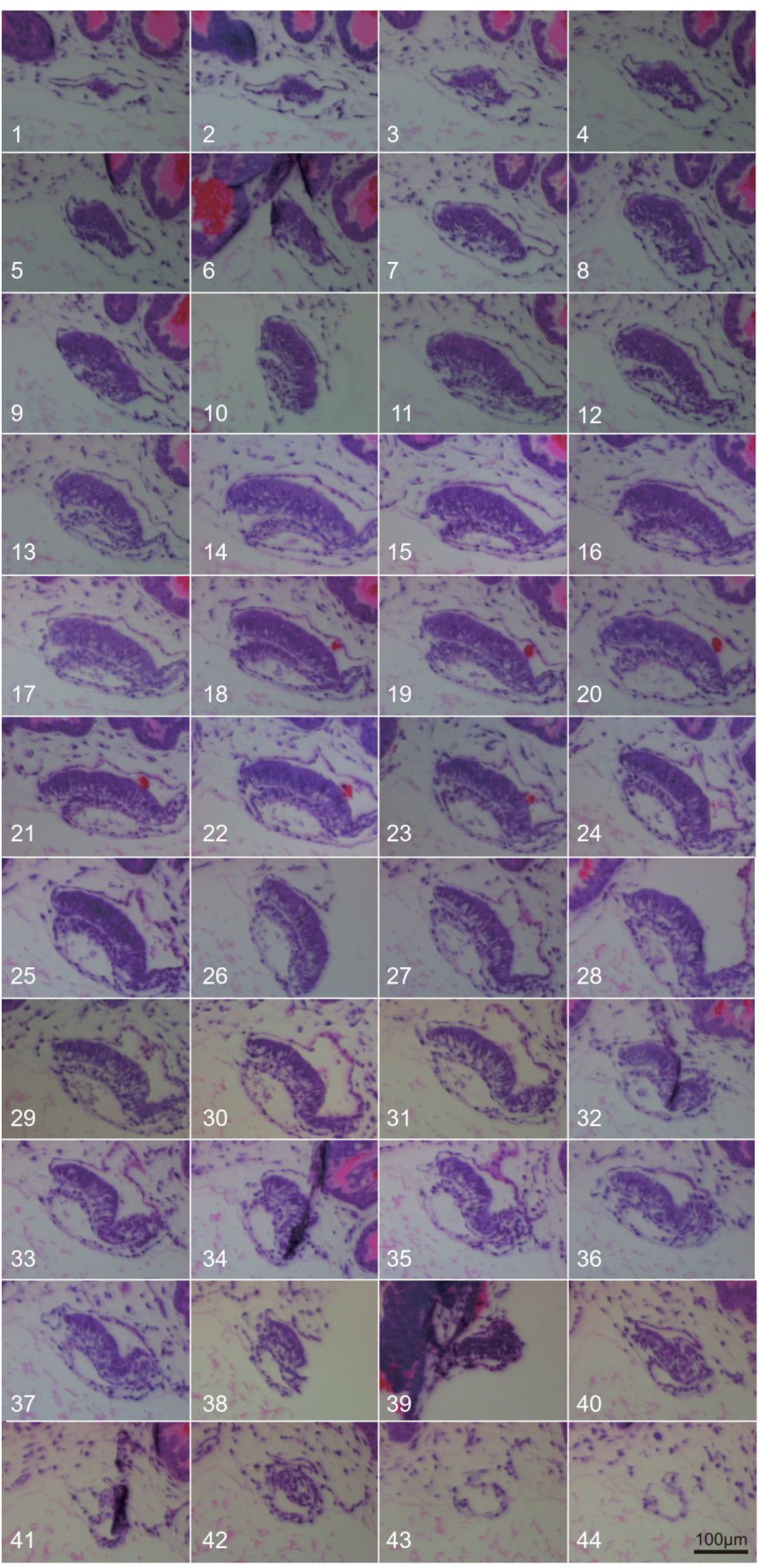
Histology of serial section of E17 monkey embryo. Haematoxylin and eosin staining of the serial sections of gastrulation embryo at E17. Scale bar, 100μm

**Figure S3.**
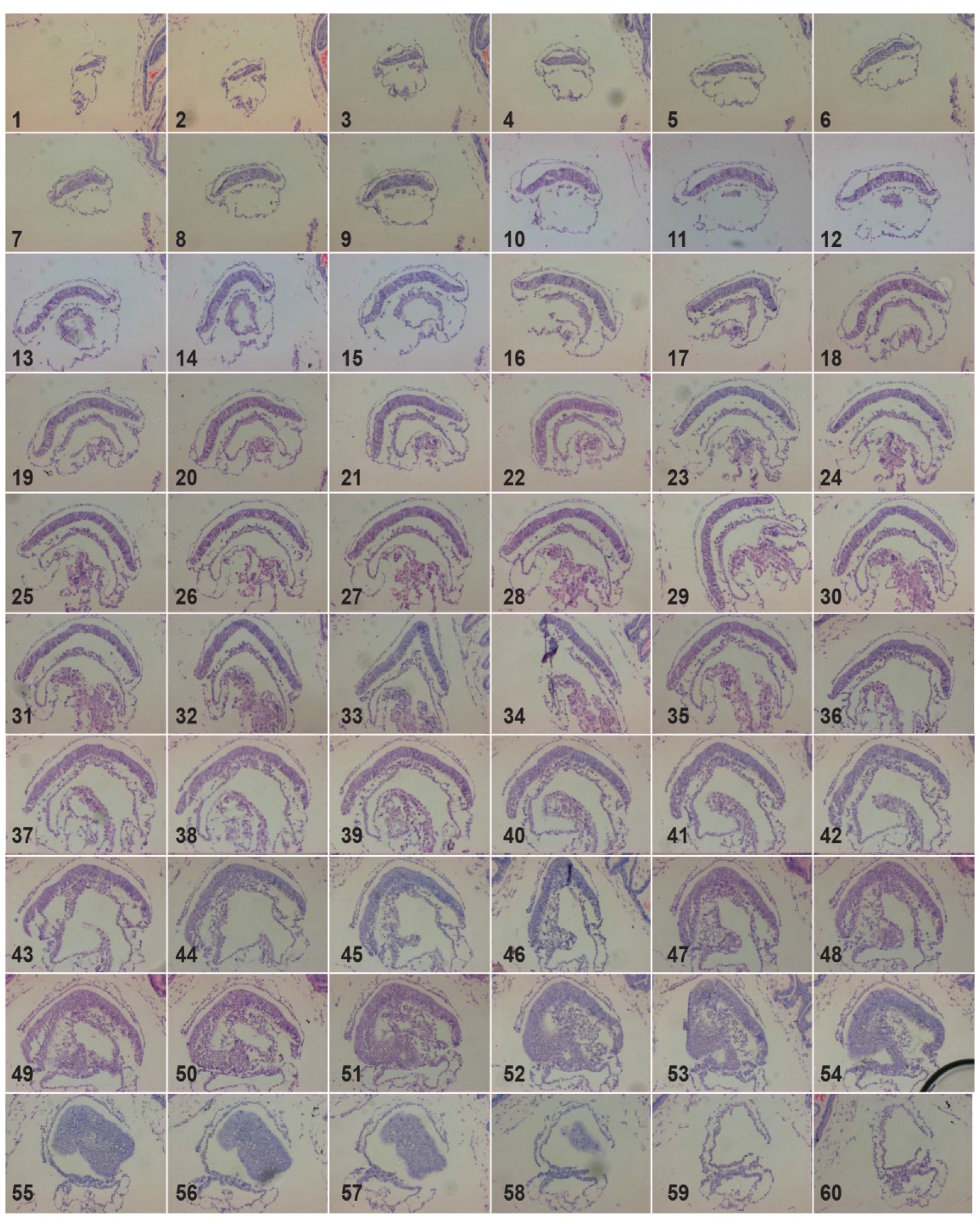
Histology of serial section of E18 monkey embryo. Haematoxylin and eosin staining of the serial sections of gastrulation embryo at E18. Scale bar, 100μm

**Figure S4.**
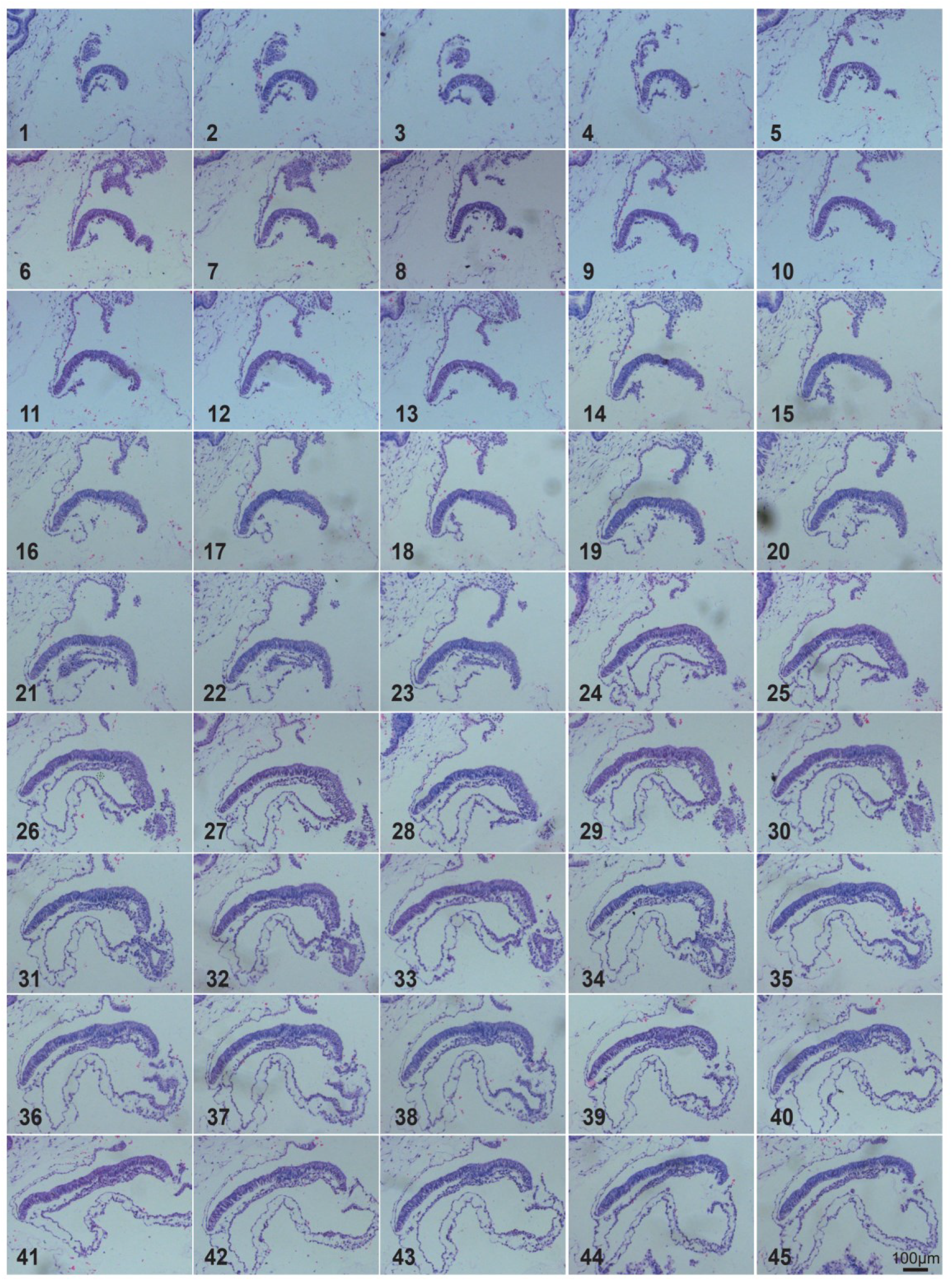
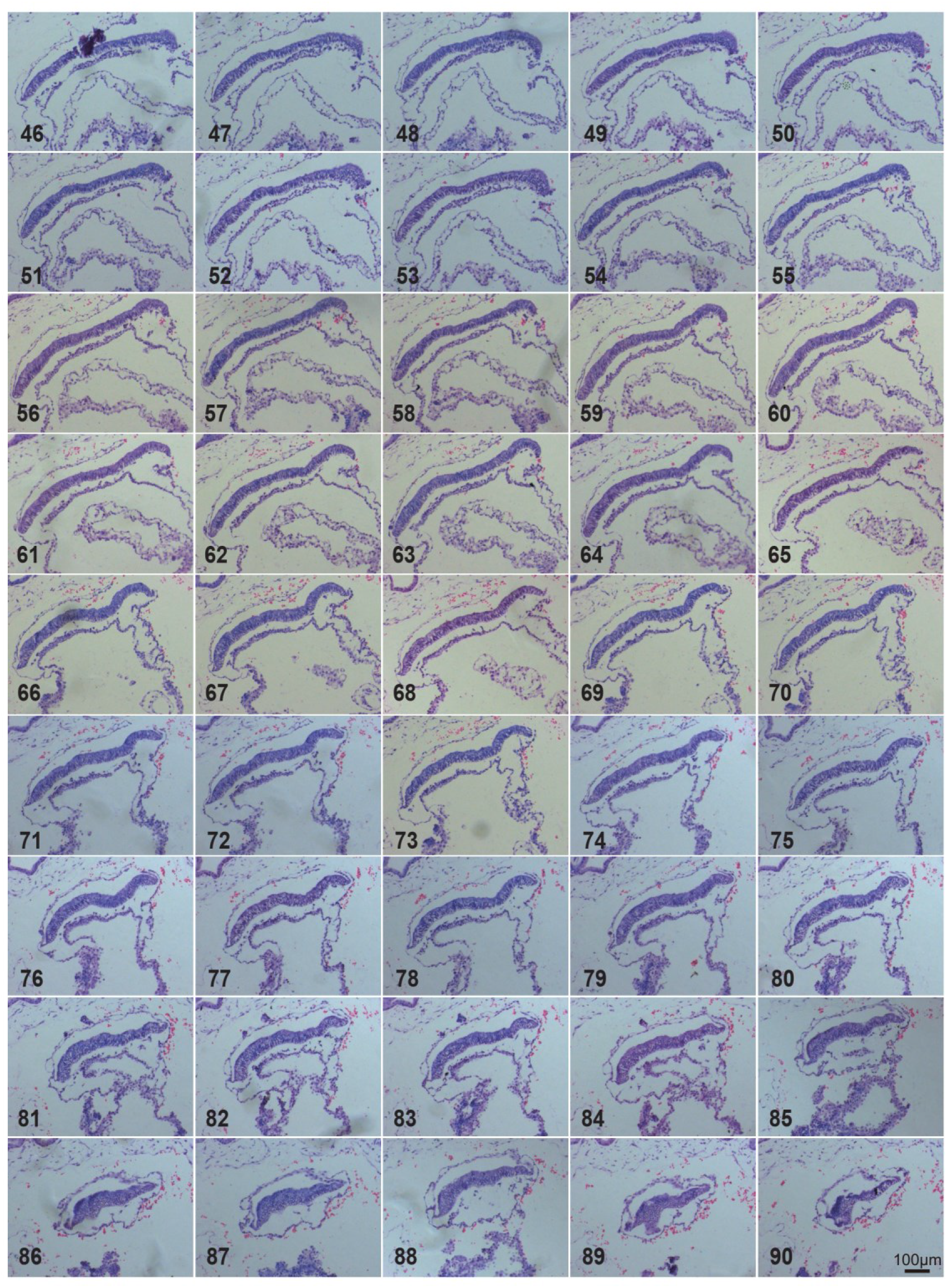
Histology of serial section of E19 monkey embryo. Haematoxylin and eosin staining of the serial sections of gastrulation embryo at E19. Scale bar, 100μm

**Figure S5.**
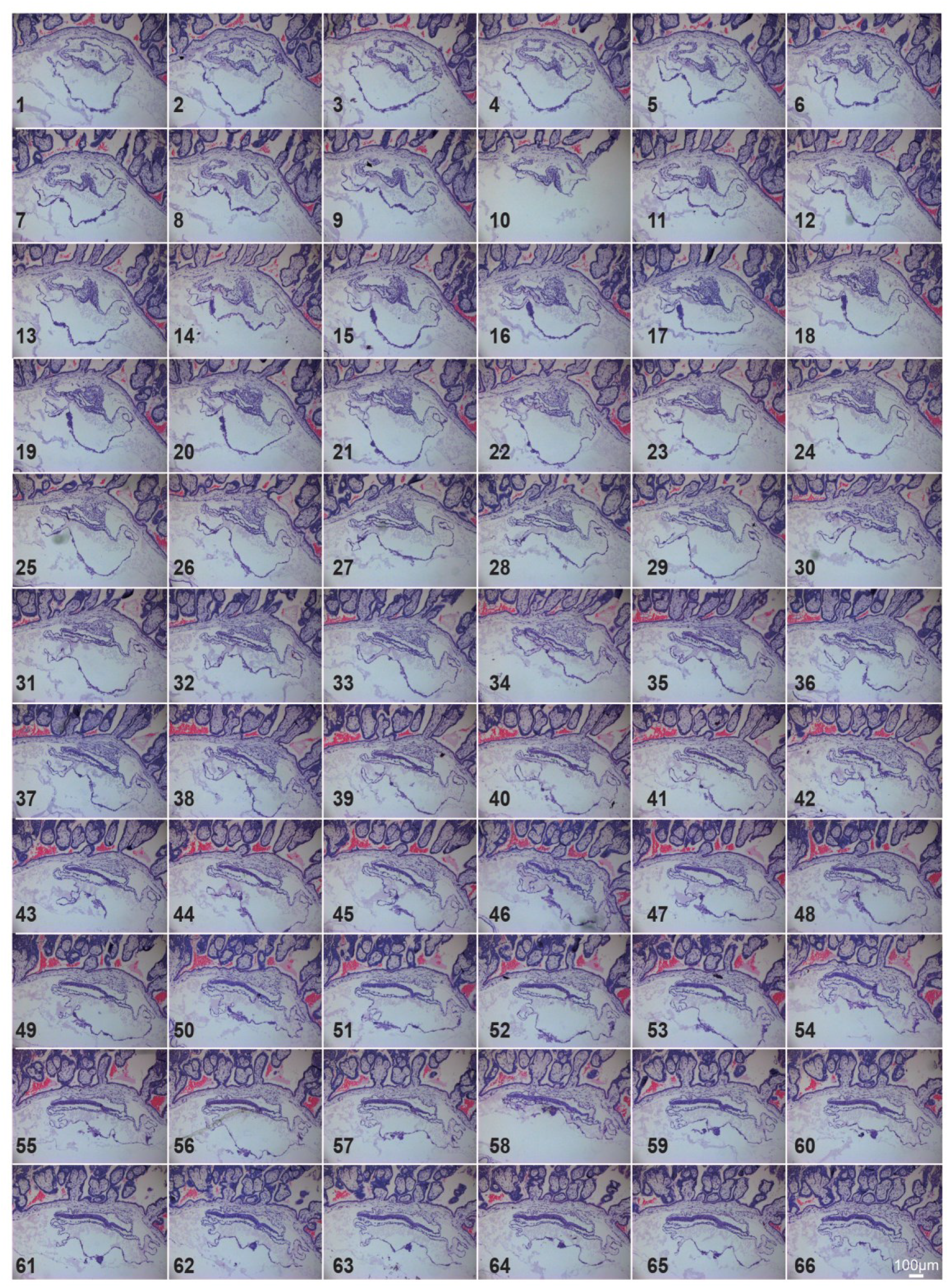
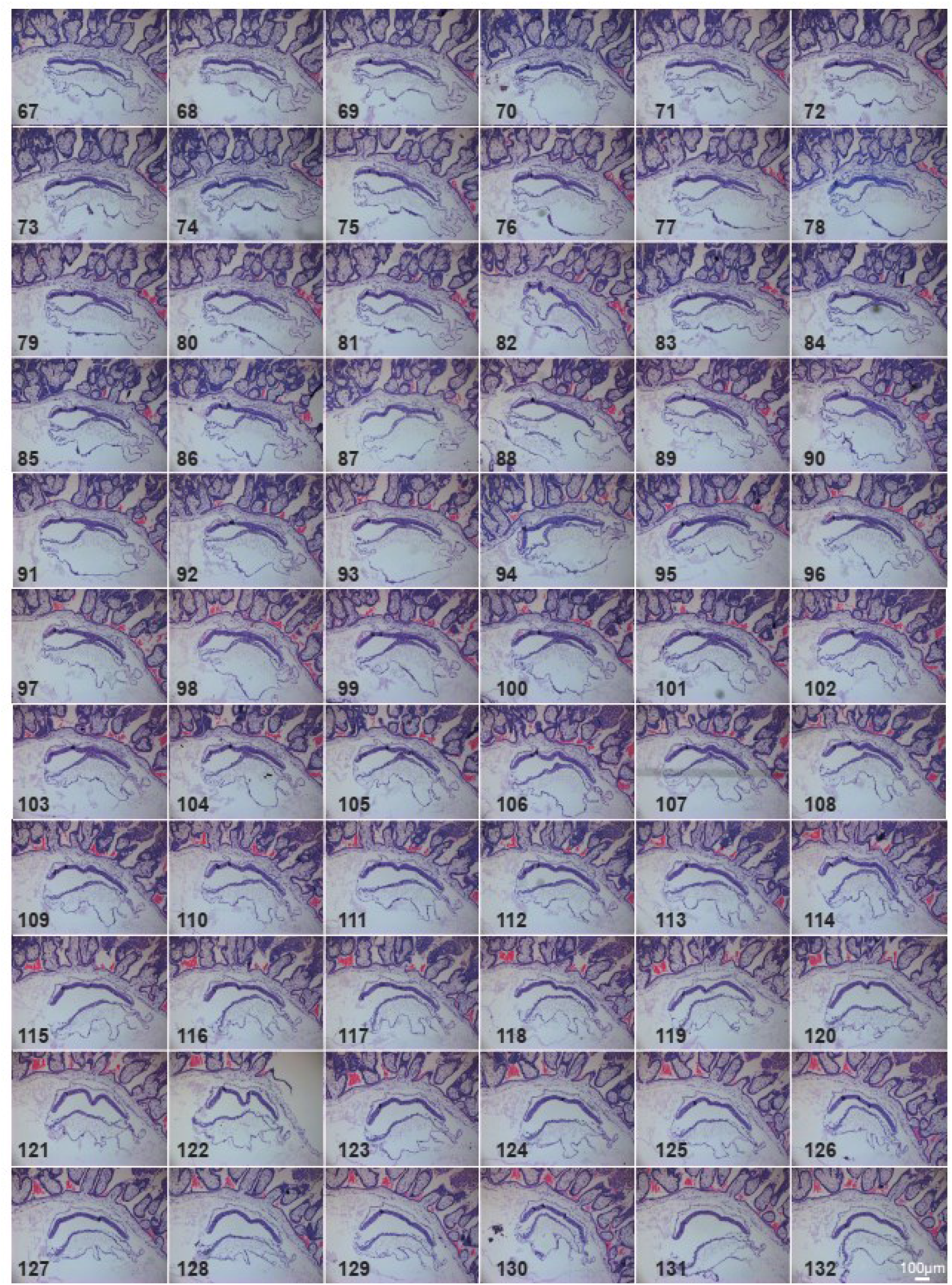
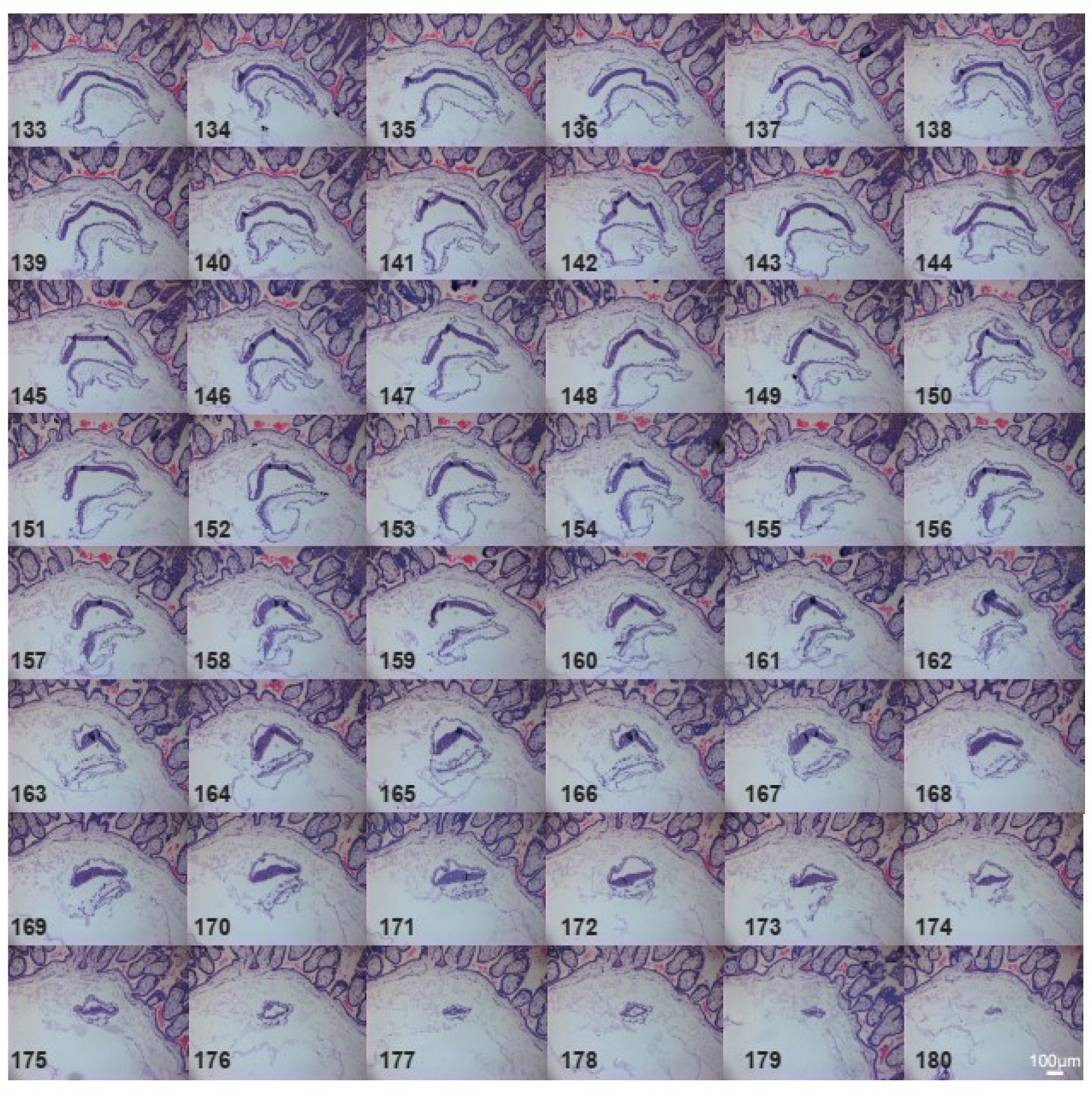
Histology of serial section of E20 monkey embryo. Haematoxylin and eosin staining of the serial sections of gastrulation embryo at E20. Scale bar, 100μm

**Figure S6.**
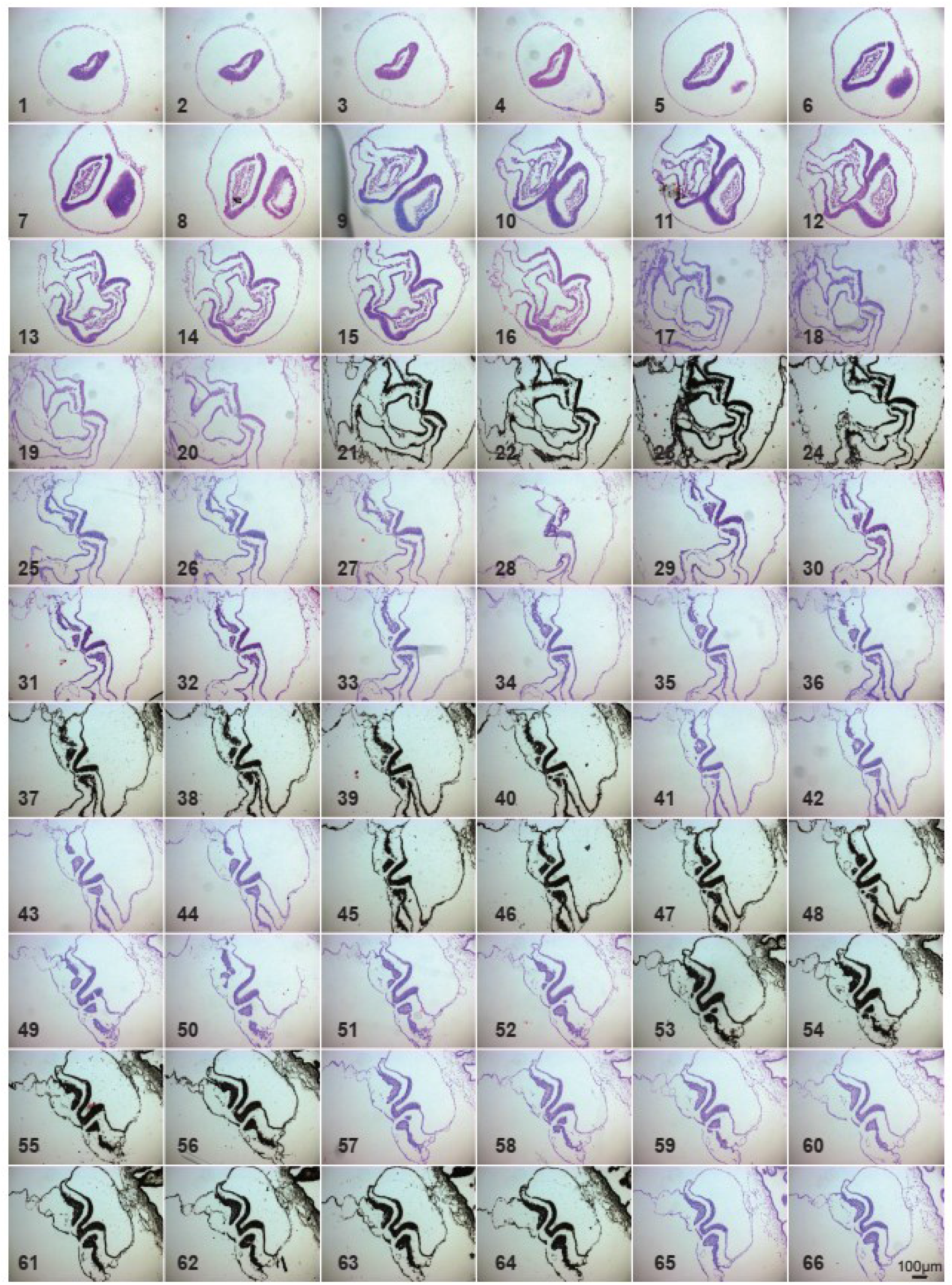
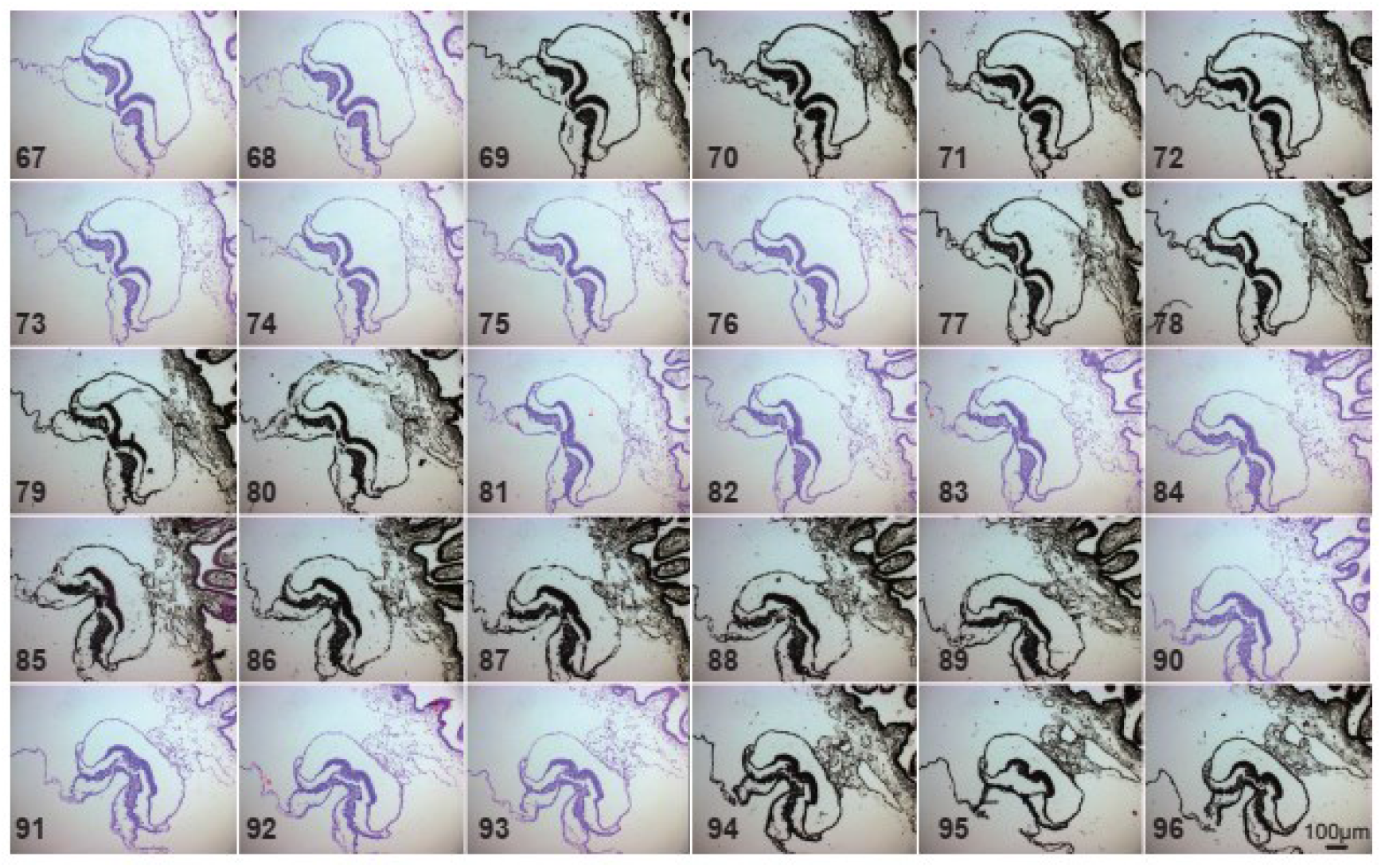
Histology of serial section of E21 monkey embryo. Haematoxylin and eosin staining of the serial sections of gastrulation embryo at E21. Scale bar, 100μm

**Figure S7.**
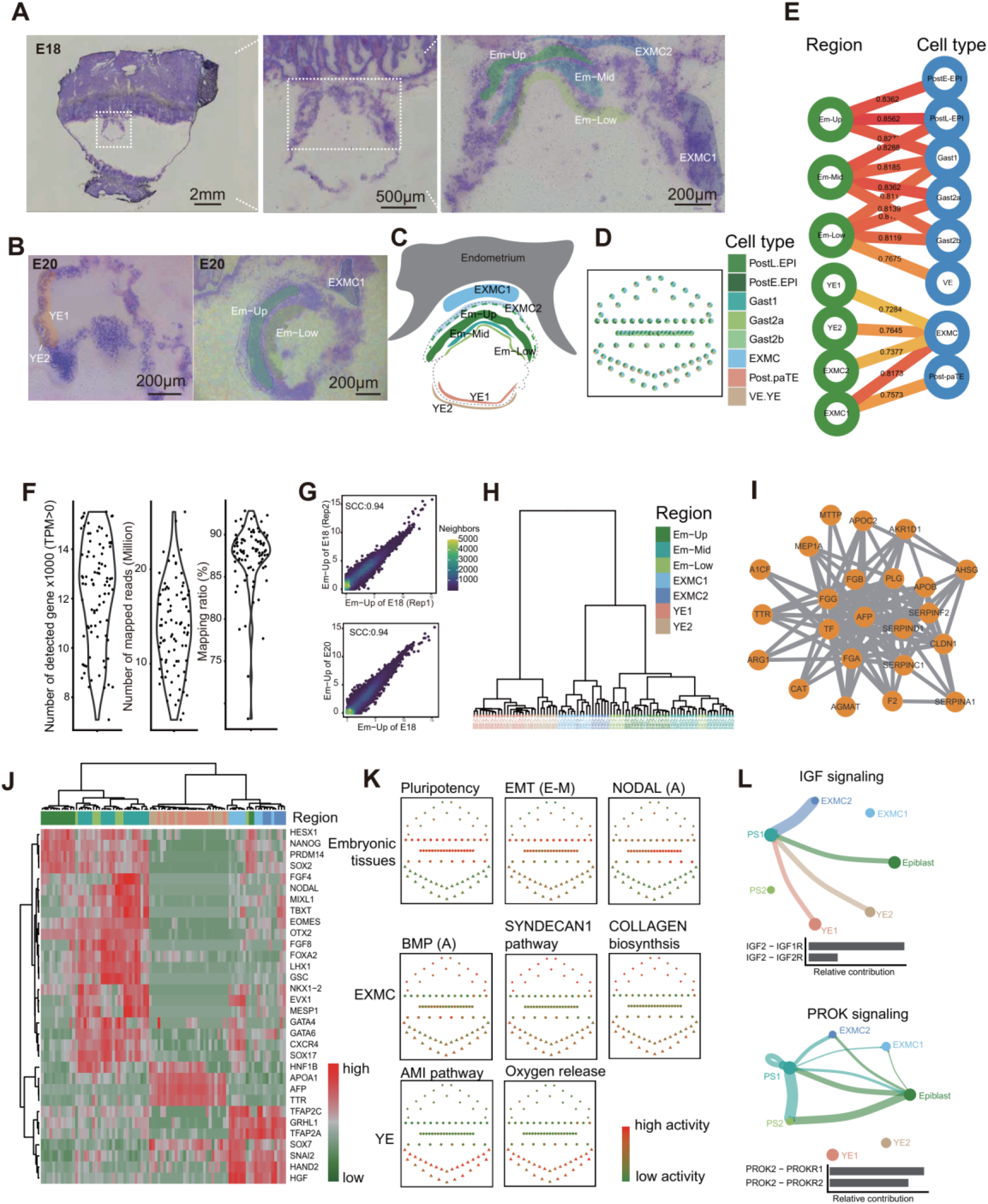
Spatial transcriptome analysis of monkey embryos at E18 and E20 by Geo-seq. A-C. Embryonic and extraembryonic tissues were captured by laser microdissection at E18 (A) and E20 (B). Representative sampling locations in 7 different regions were shaded respectively. The schematic diagram showed the collected tissue regions (C). Em-Up, upper layer of embryonic tissue; Em-Mid, middle layer of embryonic tissue; Em-Low, lower layer of embryonic tissue; EXMC1/2, part of extra-embryonic mesenchyme tissues; YE1/2, part of yolk-sac endoderm. D. Cell-type deconvolution in spatial transcriptomics samples. The single cell RNA-seq data of monkey gastrulating embryos was download from previous work GSE74767 (Nakamura et al., 2016), and the STGE-plot represents the spatial sampling in monkey embryos. The colour coding is as indicated. E. The Spearman correlation coefficient of cell population between spatial (red module) and single cell annotated cell types (blue module, download from GSE74767 (Nakamura et al., 2016)) showing the location of specific cell. The color coding is spearman correlation coefficient (SCC). F. Violin plot showing the number of detected genes (TPM > 0), number of mapped reads (Million) and mapping ratio of samples (n=86). G. The Spearman correlation coefficient (SCC) of spatial domain. Upper panel showed the SCC between embryonic replicates at E18, and lower panel presented the SCC between developmental stages. The colorbar represent the density of gene number. H. Hierarchical clustering analysis for the embryonic and extraembryonic tissues of monkey embryo based on the expression level of DEGs. Color code the different sectors. I. Weighted gene co-expression network analysis (WGCNA) represented as highest correlation coefficient values for yolk sac endoderm. J. Heatmap showing the expression pattern of germ-layer marker genes. Top colored bars indicate the sample regions as F. K. Pathway activity and epithelial-mesenchymal transition (EMT) scores analysis of the embryonic, extra-embryonic mesenchyme and yolk-sac endoderm tissues. The higher EMT scores (E-M), the more epidermal-like state. The colour coding is as indicated. L. The inferred IGF and PROK signaling networks and relative contribution of each ligand-receptor pair to those overall signaling networks. Circle sizes are proportional to the number of cells in each cell group and edge width represents the communication probability.

**Figure S8.**
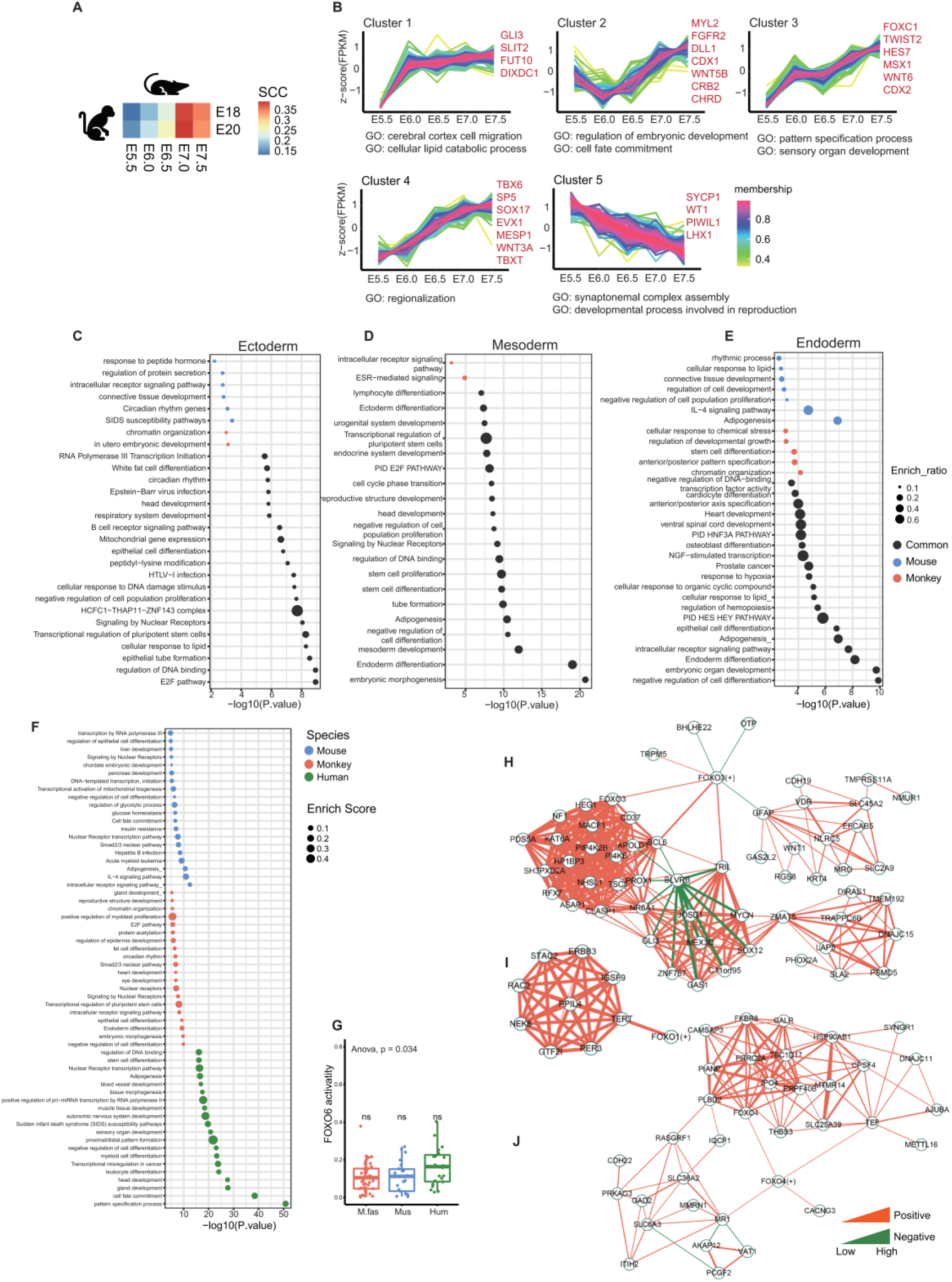
Cross-species spatial transcriptomic analysis reveals gastrulation developmental differences among mice, monkeys and humans. A, Heatmap of Spearman correlation coefficient of expressed gene between mouse and monkey gastrulating embryos. B, Cluster analysis of the gene expression patterns across the gastrulation development stages. C-E, Dotplot showing functional enrichment of species – specificity regulons in ectoderm (C), mesoderm (D) and endoderm (E) formation between mouse and macaque. F, Functional enrichment, interactome analysis and gene annotation based on species – specificity regulons. G, The activities of FOXO6 regulons between mouse, monkey and human gastrulation embryonic tissue. H-J, Networks showing the relationship of FOXO1 (H), FOXO3 (I) and FOXO4 (J) with their targets and top 10 related genes, respectively. The color of edges means positive interaction (orange) or negative interaction (green), and the width of edges represents the strength of the correlation.

**Table S1. List of the length and thickness of sectioned epiblast from E17 to E20, Related to Figure 1**

The gastrulating embryo sizes were assessed by the length and thickness of sectioned epiblast.

**Table S2. Cell counting of germ layers in gastrulating Cynomolgus monkey embryos, Related to Figure 1**

The detailed histological analysis was applied to determine cell number by counting nuclei on every section of the gastrulating embryos. The total score was then adjusted by applying Abercrombie’s correction formula to give an estimate of the actual cell number.

**Table S3. The interspecies differences underlying the germ layer segregation, Related to Figure 7**

The species/spatial domain-specificity score (SSS) of mouse and macaque in the process of ectoderm, mesoderm and endoderm specification was showed in sheet 1. The SSS and p-value of interspecies differences in mouse, macaque and human were listed in sheet 2. The species-specific regulons were listed in sheet 3.

## REFERENCE

Abe, T., Sakaue-Sawano, A., Kiyonari, H., Shioi, G., Inoue, K., Horiuchi, T., Nakao, K., Miyawaki, A., Aizawa, S., and Fujimori, T. (2013). Visualization of cell cycle in mouse embryos with Fucci2 reporter directed by Rosa26 promoter. Development 140, 237–246.

Abercrombie, M. (1946). Estimation of nuclear population from microtome sections. The Anatomical record 94, 239–247.

Aibar, S., Gonzalez-Blas, C.B., Moerman, T., Huynh-Thu, V.A., Imrichova, H., Hulselmans, G., Rambow, F., Marine, J.C., Geurts, P., Aerts, J., et al. (2017). SCENIC: single-cell regulatory network inference and clustering. Nature methods.

Andreatta, M., and Carmona, S.J. (2021). UCell: Robust and scalable single-cell gene signature scoring. Computational and structural biotechnology journal 19, 3796–3798.

Baardman, M.E., Kerstjens-Frederikse, W.S., Berger, R.M., Bakker, M.K., Hofstra, R.M., and Plösch, T. (2013). The role of maternal-fetal cholesterol transport in early fetal life: current insights. Biology of reproduction 88, 24.

Ben-Haim, N., Lu, C., Guzman-Ayala, M., Pescatore, L., Mesnard, D., Bischofberger, M., Naef, F., Robertson, E.J., and Constam, D.B. (2006). The nodal precursor acting via activin receptors induces mesoderm by maintaining a source of its convertases and BMP4. Developmental cell 11, 313–323.

Bianchi, D.W., Wilkins-Haug, L.E., Enders, A.C., and Hay, E.D. (1993). Origin of extraembryonic mesoderm in experimental animals: relevance to chorionic mosaicism in humans. American journal of medical genetics 46, 542–550.

Boroviak, T., Stirparo, G.G., Dietmann, S., Hernando-Herraez, I., Mohammed, H., Reik, W., Smith, A., Sasaki, E., Nichols, J., and Bertone, P. (2018). Single cell transcriptome analysis of human, marmoset and mouse embryos reveals common and divergent features of preimplantation development. Development 145.

Brennan, J., Lu, C.C., Norris, D.P., Rodriguez, T.A., Beddington, R.S., and Robertson, E.J. (2001). Nodal signalling in the epiblast patterns the early mouse embryo. Nature 411, 965–969.

Bu, D., Luo, H., Huo, P., Wang, Z., Zhang, S., He, Z., Wu, Y., Zhao, L., Liu, J., Guo, J., et al. (2021). KOBAS-i: intelligent prioritization and exploratory visualization of biological functions for gene enrichment analysis. Nucleic acids research 49, W317–W325.

Carlson, B.M. (2015). Gastrulation and Germ Layer Formation. In Reference Module in Biomedical Sciences.

Chakraborty, P., George, J.T., Tripathi, S., Levine, H., and Jolly, M.K. (2020). Comparative Study of Transcriptomics-Based Scoring Metrics for the Epithelial-Hybrid-Mesenchymal Spectrum. Front Bioeng Biotechnol 8, 220.

Chakravarti, D., LaBella, K.A., and DePinho, R.A. (2021). Telomeres: history, health, and hallmarks of aging. Cell 184, 306–322.

Chen, J., Suo, S., Tam, P.P., Han, J.J., Peng, G., and Jing, N. (2017). Spatial transcriptomic analysis of cryosectioned tissue samples with Geo-seq. Nature protocols 12, 566–580.

Chen, S., Zhou, Y., Chen, Y., and Gu, J. (2018). fastp: an ultra-fast all-in-one FASTQ preprocessor. Bioinformatics 34, i884–i890.

Chhabra, S., Liu, L., Goh, R., Kong, X., and Warmflash, A. (2019). Dissecting the dynamics of signaling events in the BMP, WNT, and NODAL cascade during self-organized fate patterning in human gastruloids. PLoS biology 17, e3000498.

Cindrova-Davies, T., Jauniaux, E., Elliot, M.G., Gong, S., Burton, G.J., and Charnock-Jones, D.S. (2017). RNA-seq reveals conservation of function among the yolk sacs of human, mouse, and chicken. Proceedings of the National Academy of Sciences of the United States of America 114, E4753–E4761.

Ciruna, B., and Rossant, J. (2001). FGF Signaling Regulates Mesoderm Cell Fate Specification and Morphogenetic Movement at the Primitive Streak. Developmental cell 1, 37–49.

Cui, G., Chen, J., Chen, S., Qian, Y., Peng, G., and Jing, N. (2019). Spatio-temporal transcriptome construction of early mouse embryo with Geo-seq and Auto-seq. Protocol Exchange, 10.21203/rs.21202.10081/v21201.

de Bakker, B.S., de Jong, K.H., Hagoort, J., de Bree, K., Besselink, C.T., de Kanter, F.E.C., Veldhuis, T., Bais, B., Schildmeijer, R., Ruijter, J.M., et al. (2016). An interactive three-dimensional digital atlas and quantitative database of human development. Science 354, aag0053.

DeTomaso, D., Jones, M.G., Subramaniam, M., Ashuach, T., Ye, C.J., and Yosef, N. (2019). Functional interpretation of single cell similarity maps. Nature communications 10, 4376.

Dong, D., and Yang, P. (2018). Yolk Sac. In Encyclopedia of Reproduction (Second Edition), M.K. Skinner, ed. (Oxford: Academic Press), pp. 551–558.

Dongre, A., and Weinberg, R.A. (2019). New insights into the mechanisms of epithelial-mesenchymal transition and implications for cancer. Nature reviews Molecular cell biology 20, 69–84.

Drost, H.-G. (2018). Philentropy: Information Theory and Distance Quantification with R. Journal of Open Source Software 3.

Dufort, D., Schwartz, L., Harpal, K., and Rossant, J. (1998). The transcription factor HNF3beta is required in visceral endoderm for normal primitive streak morphogenesis. Development 125, 3015–3025.

Eivers, E., McCarthy, K., Glynn, C., Nolan, C.M., and Byrnes, L. (2004). Insulin-like growth factor (IGF) signalling is required for early dorso-anterior development of the zebrafish embryo. The International journal of developmental biology 48, 1131–1140.

Elosua-Bayes, M., Nieto, P., Mereu, E., Gut, I., and Heyn, H. (2021). SPOTlight: seeded NMF regression to deconvolute spatial transcriptomics spots with single-cell transcriptomes. Nucleic acids research.

Enders, A.C., and King, B.F. (1988). Formation and differentiation of extraembryonic mesoderm in the rhesus monkey. The American journal of anatomy 181, 327–340.

Enders, A.C., Schlafke, S., and Hendrickx, A.G. (1986). Differentiation of the embryonic disc, amnion, and yolk sac in the rhesus monkey. The American journal of anatomy 177, 161–185.

Exalto, N. (1995). Early human nutrition. European Journal of Obstetrics & Gynecology and Reproductive Biology 61, 3–6.

Feng, S., Xing, C., Shen, T., Qiao, Y., Wang, R., Chen, J., Liao, J., Lu, Z., Yang, X., Abd-Allah, S.M., et al. (2017). Abnormal Paraventricular Nucleus of Hypothalamus and Growth Retardation Associated with Loss of Nuclear Receptor Gene COUP-TFII. Scientific reports 7, 5282.

Fu, J., Warmflash, A., and Lutolf, M.P. (2020). Stem-cell-based embryo models for fundamental research and translation. Nature materials.

Gao, Y., Cao, Q., Lu, L., Zhang, X., Zhang, Z., Dong, X., Jia, W., and Cao, Y. (2015). Kruppel-like factor family genes are expressed during Xenopus embryogenesis and involved in germ layer formation and body axis patterning. 244, 1328–1346.

Ghimire, S., Mantziou, V., Moris, N., and Arias, A.M. (2021). Human gastrulation: The embryo and its models. Developmental biology.

Grapin-Botton, A., and Constam, D. (2007). Evolution of the mechanisms and molecular control of endoderm formation. Mechanisms of development 124, 253–278.

Grobstein, C. (1985). The early development of human embryos. The Journal of medicine and philosophy 10, 213–236.

Gualdi, R., Bossard, P., Zheng, M., Hamada, Y., Coleman, J.R., and Zaret, K.S. (1996). Hepatic specification of the gut endoderm in vitro: cell signaling and transcriptional control. Genes & development 10, 1670–1682.

Hannenhalli, S., and Kaestner, K.H. (2009). The evolution of Fox genes and their role in development and disease. Nature reviews Genetics 10, 233–240.

Hendrickx, A.G. (1972). Early development of the embryo in non-human primates and man. Acta endocrinologica Supplementum 166, 103–130.

Hyun, I., Bredenoord, A.L., Briscoe, J., Klipstein, S., and Tan, T. (2021). Human embryo research beyond the primitive streak. Science 371, 998–1000.

Jin, S., Guerrero-Juarez, C.F., Zhang, L., Chang, I., Ramos, R., Kuan, C.H., Myung, P., Plikus, M.V., and Nie, Q. (2021). Inference and analysis of cell-cell communication using CellChat. Nature communications 12, 1088.

Kaestner, K.H., Hiemisch, H., and Schütz, G. (1998). Targeted disruption of the gene encoding hepatocyte nuclear factor 3gamma results in reduced transcription of hepatocyte-specific genes. Molecular and cellular biology 18, 4245–4251.

Lai, E., Prezioso, V.R., Tao, W.F., Chen, W.S., and Darnell, J.E., Jr. (1991). Hepatocyte nuclear factor 3 alpha belongs to a gene family in mammals that is homologous to the Drosophila homeotic gene fork head. Genes & development 5, 416–427.

Lamouille, S., Xu, J., and Derynck, R. (2014). Molecular mechanisms of epithelial-mesenchymal transition. Nature reviews Molecular cell biology 15, 178–196.

Langfelder, P., and Horvath, S. (2008). WGCNA: an R package for weighted correlation network analysis. BMC bioinformatics 9, 559.

Larsson, L., Frisen, J., and Lundeberg, J. (2021). Spatially resolved transcriptomics adds a new dimension to genomics. Nature methods 18, 15–18.

Lawson, K.A., and Wilson, V. (2016). A Revised Staging of Mouse Development Before Organogenesis. In Kaufman’s Atlas of Mouse Development Supplement, pp. 51–64.

Luckett, W.P. (1978). Origin and differentiation of the yolk sac and extraembryonic mesoderm in presomite human and rhesus monkey embryos. The American journal of anatomy 152, 59–97.

Ma, H., Zhai, J., Wan, H., Jiang, X., Wang, X., Wang, L., Xiang, Y., He, X., Zhao, Z.A., Zhao, B., et al. (2019). In vitro culture of Cynomolgus monkey embryos beyond early gastrulation. Science 366.

Massri, A.J., Schiebinger, G.R., Berrio, A., Wang, L., Wray, G.A., and McClay, D.R. (2021). The Epithelialto Mesenchymal Transition. Methods Mol Biol 2179, 303–314.

McDole, K., Guignard, L., Amat, F., Berger, A., Malandain, G., Royer, L.A., Turaga, S.C., Branson, K., and Keller, P.J. (2018). In Toto Imaging and Reconstruction of Post-Implantation Mouse Development at the Single-Cell Level. Cell.

Mengjun, L.G. (2019). TCseq: Time course sequencing data analysis.

Mitiku, N., and Baker, J.C. (2007). Genomic analysis of gastrulation and organogenesis in the mouse. Developmental cell 13, 897–907.

Mittnenzweig, M., Mayshar, Y., Cheng, S., Ben-Yair, R., Hadas, R., Rais, Y., Chomsky, E., Reines, N., Uzonyi, A., Lumerman, L., et al. (2021). A single-embryo, single-cell time-resolved model for mouse gastrulation. Cell.

Monaghan, A.P., Kaestner, K.H., Grau, E., and Schütz, G. (1993). Postimplantation expression patterns indicate a role for the mouse forkhead/HNF-3 alpha, beta and gamma genes in determination of the definitive endoderm, chordamesoderm and neuroectoderm. Development 1 *9*, 567–578.

Moore, H.D., Gems, S., and Hearn, J.P. (1985). Early implantation stages in the marmoset monkey (Callithrix jacchus). The American journal of anatomy 172, 265–278.

Moris, N., Anlas, K., van den Brink, S.C., Alemany, A., Schröder, J., Ghimire, S., Balayo, T., van Oudenaarden, A., and Martinez Arias, A. (2020). An in vitro model of early anteroposterior organization during human development. Nature 582, 410–415.

Muller, F., and O’Rahilly, R. (2004). The primitive streak, the caudal eminence and related structures in staged human embryos. Cells, tissues, organs 177, 2–20.

Nakamura, T., Fujiwara, K., Saitou, M., and Tsukiyama, T. (2021). Non-human primates as a model for human development. Stem Cell Reports 16, 1093–1103.

Nakamura, T., Okamoto, I., Sasaki, K., Yabuta, Y., Iwatani, C., Tsuchiya, H., Seita, Y., Nakamura, S., Yamamoto, T., and Saitou, M. (2016). A developmental coordinate of pluripotency among mice, monkeys and humans. Nature 537, 57–62.

Nicetto, D., Donahue, G., Jain, T., Peng, T., Sidoli, S., Sheng, L., Montavon, T., Becker, J.S., Grindheim, J.M., Blahnik, K., et al. (2019). H3K9me3-heterochromatin loss at protein-coding genes enables developmental lineage specification. 363, 294–297.

Nikolopoulou, E., Galea, G.L., Rolo, A., Greene, N.D., and Copp, A.J. (2017). Neural tube closure: cellular, molecular and biomechanical mechanisms. Development 144, 552–566.

Niu, Y., Sun, N., Li, C., Lei, Y., Huang, Z., Wu, J., Si, C., Dai, X., Liu, C., Wei, J., et al. (2019). Dissecting primate early post-implantation development using long-term in vitro embryo culture. Science 366, 5754.

Nowotschin, S., Hadjantonakis, A.K., and Campbell, K. (2019). The endoderm: a divergent cell lineage with many commonalities. Development 146, dev150920.

O’Rahilly, R., and Muller, F. (2010). Developmental stages in human embryos: revised and new measurements. Cells, tissues, organs 192, 73–84.

Otasek, D., Morris, J.H., Boucas, J., Pico, A.R., and Demchak, B. (2019). Cytoscape Automation: empowering workflow-based network analysis. Genome biology 20, 185.

Peng, G., Cui, G., Ke, J., and Jing, N. (2020a). Using Single-Cell and Spatial Transcriptomes to Understand Stem Cell Lineage Specification During Early Embryo Development. Annual Review of Genomics and Human Genetics 21, 163–181.

Peng, G., Cui, G., Ke, J., and Jing, N. (2020b). Using Single-Cell and Spatial Transcriptomes to Understand Stem Cell Lineage Specification During Early Embryo Development. Annu Rev Genomics Hum Genet 21, 163–181.

Peng, G., Suo, S., Chen, J., Chen, W., Liu, C., Yu, F., Wang, R., Chen, S., Sun, N., Cui, G., et al. (2016). Spatial Transcriptome for the Molecular Annotation of Lineage Fates and Cell Identity in Mid-gastrula Mouse Embryo. Developmental cell 36, 681–697.

Peng, G., Suo, S., Cui, G., Yu, F., Wang, R., Chen, J., Chen, S., Liu, Z., Chen, G., Qian, Y., et al. (2019). Molecular architecture of lineage allocation and tissue organization in early mouse embryo. Nature 572, 528–532.

Peng, H., Ruan, Z., Long, F., Simpson, J.H., and Myers, E.W. (2010). V3D enables real-time 3D visualization and quantitative analysis of large-scale biological image data sets. Nat Biotechnol 28, 348–353.

Pertea, M., Kim, D., Pertea, G.M., Leek, J.T., and Salzberg, S.L. (2016). Transcript-level expression analysis of RNA-seq experiments with HISAT, StringTie and Ballgown. Nature protocols 11, 1650–1667.

Pfister, S., Steiner, K.A., and Tam, P.P. (2007). Gene expression pattern and progression of embryogenesis in the immediate post-implantation period of mouse development. Gene expression patterns : GEP 7, 558–573.

Pijuan-Sala, B., Guibentif, C., and Gottgens, B. (2018). Single-cell transcriptional profiling: a window into embryonic cell-type specification. Nature reviews Molecular cell biology.

Rivera-Perez, J.A., and Hadjantonakis, A.K. (2015). The Dynamics of Morphogenesis in the Early Mouse Embryo. Cold Spring Harb Perspect Biol 7.

Rivera-Perez, J.A., and Magnuson, T. (2005). Primitive streak formation in mice is preceded by localized activation of Brachyury and Wnt3. Developmental biology 288, 363–371.

Ross, C., and Boroviak, T.E. (2020). Origin and function of the yolk sac in primate embryogenesis. Nature communications 11, 3760.

Sasaki, H., and Hogan, B.L. (1993). Differential expression of multiple fork head related genes during gastrulation and axial pattern formation in the mouse embryo. Development 118, 47–59.

Scheibner, K., Schirge, S., Burtscher, I., Buttner, M., Sterr, M., Yang, D., Bottcher, A., Ansarullah, Irmler, M., Beckers, J., et al. (2021). Epithelial cell plasticity drives endoderm formation during gastrulation. Nat Cell Biol.

Schindelin, J., Arganda-Carreras, I., Frise, E., Kaynig, V., Longair, M., Pietzsch, T., Preibisch, S., Rueden, C., Saalfeld, S., Schmid, B., et al. (2012). Fiji: an open-source platform for biological-image analysis. Nature methods 9, 676–682.

Shahbazi, M.N. (2020). Mechanisms of human embryo development: from cell fate to tissue shape and back. Development 147, dev190629.

Sheaffer, K.L., and Kaestner, K.H. (2012). Transcriptional networks in liver and intestinal development. Cold Spring Harb Perspect Biol 4, a008284.

Singh, M., Yelle, N., Venugopal, C., and Singh, S.K. (2018). EMT: Mechanisms and therapeutic implications. Pharmacol Ther 182, 80–94.

Snow, M. (1977). Gastrulation in the mouse: growth and regionalization of the epiblast. Journal of embryology and experimental morphology 42, 293–303.

Souilhol, C., Perea-Gomez, A., Camus, A., Beck-Cormier, S., Vandormael-Pournin, S., Escande, M., Collignon, J., and Cohen-Tannoudji, M. (2015). NOTCH activation interferes with cell fate specification in the gastrulating mouse embryo. Development 142, 3649–3660.

Stuart, T., Butler, A., Hoffman, P., Hafemeister, C., Papalexi, E., Mauck, W.M., 3rd, Hao, Y., Stoeckius, M., Smibert, P., and Satija, R. (2019). Comprehensive Integration of Single-Cell Data. Cell 177, 1888–1902 e1821.

Sun, X., Meyers, E.N., Lewandoski, M., and Martin, G.R. (1999). Targeted disruption of Fgf8 causes failure of cell migration in the gastrulating mouse embryo. Genes & development 13, 1834–1846.

Suo, S., Zhu, Q., Saadatpour, A., Fei, L., Guo, G., and Yuan, G.C. (2018). Revealing the Critical Regulators of Cell Identity in the Mouse Cell Atlas. Cell Rep 25, 1436–1445 e1433.

Tam, P.P., and Behringer, R.R. (1997). Mouse gastrulation: the formation of a mammalian body plan. Mechanisms of development 68, 3–25.

Tam, P.P.L., and Ho, J.W.K. (2020). Cellular diversity and lineage trajectory: insights from mouse single cell transcriptomes. Development 147.

Tang, T.T.-L., Dowbenko, D., Jackson, A., Toney, L., Lewin, D.A., Dent, A.L., and Lasky, L.A. (2002). The Forkhead Transcription Factor AFX Activates Apoptosis by Induction of the BCL-6 Transcriptional Repressor *. Journal of Biological Chemistry 277, 14255–14265.

Tani, S., Chung, U.I., Ohba, S., and Hojo, H. (2020). Understanding paraxial mesoderm development and sclerotome specification for skeletal repair. Exp Mol Med.

Tarara, R., Enders, A.C., Hendrickx, A.G., Gulamhusein, N., Hodges, J.K., Hearn, J.P., Eley, R.B., and Else, J.G. (1987). Early implantation and embryonic development of the baboon: stages 5, 6 and 7. Anat Embryol (Berl) 176, 267–275.

Thieme, R., Ramin, N., Fischer, S., Puschel, B., Fischer, B., and Santos, A.N. (2012). Gastrulation in rabbit blastocysts depends on insulin and insulin-like-growth-factor 1. Molecular and cellular endocrinology 348, 112–119.

Tyser, R.C.V., Mahammadov, E., Nakanoh, S., Vallier, L., Scialdone, A., and Srinivas, S. (2020). A spatially resolved single cell atlas of human gastrulation.

Tyser, R.C.V., Mahammadov, E., Nakanoh, S., Vallier, L., Scialdone, A., and Srinivas, S. (2021). Single-cell transcriptomic characterization of a gastrulating human embryo. Nature.

Varelas, X., Sakuma, R., Samavarchi-Tehrani, P., Peerani, R., Rao, B.M., Dembowy, J., Yaffe, M.B., Zandstra, P.W., and Wrana, J.L. (2008). TAZ controls Smad nucleocytoplasmic shuttling and regulates human embryonic stem-cell self-renewal. Nat Cell Biol 10, 837–848.

Viotti, M., Nowotschin, S., and Hadjantonakis, A.K. (2014). SOX17 links gut endoderm morphogenesis and germ layer segregation. Nat Cell Biol 16, 1146–1156.

Wang, C., Liu, X., Gao, Y., Yang, L., Li, C., Liu, W., Chen, C., Kou, X., Zhao, Y., Chen, J., et al. (2018). Reprogramming of H3K9me3-dependent heterochromatin during mammalian embryo development. Nat Cell Biol 20, 620–631.

Wong, E.A., and Uni, Z. (2021). Centennial Review: The chicken yolk sac is a multifunctional organ. Poult Sci 100, 100821.

Xiang, L., Yin, Y., Zheng, Y., Ma, Y., Li, Y., Zhao, Z., Guo, J., Ai, Z., Niu, Y., Duan, K., et al. (2019). A developmental landscape of 3D-cultured human pre-gastrulation embryos. Nature 577, 537–542.

Yamasaki, J., Iwatani, C., Tsuchiya, H., Okahara, J., Sankai, T., and Torii, R. (2011). Vitrification and transfer of Cynomolgus monkey (Macaca fascicularis) embryos fertilized by intracytoplasmic sperm injection. Theriogenology 76, 33–38.

Yamashita, S., Ogawa, K., Ikei, T., Fujiki, T., and Katakura, Y. (2014). FOXO3a potentiates hTERT gene expression by activating c-MYC and extends the replicative life-span of human fibroblast. PloS one 9, e101864.

Yang, J., Wang, B., Chen, H., Chen, X., Li, J., Chen, Y., Yuan, D., and Zheng, S. (2019). Thyrotroph embryonic factor is downregulated in bladder cancer and suppresses proliferation and tumorigenesis via the AKT/FOXOs signalling pathway. Cell Prolif 52, e12560.

Yang, R., Goedel, A., Kang, Y., Si, C., Chu, C., Zheng, Y., Chen, Z., Gruber, P.J., Xiao, Y., Zhou, C., et al. (2021). Amnion signals are essential for mesoderm formation in primates. Nature communications 12, 5126.

Yeung, T.L., Leung, C.S., Wong, K.K., Gutierrez-Hartmann, A., Kwong, J., Gershenson, D.M., and Mok, S.C. (2017). ELF3 is a negative regulator of epithelial-mesenchymal transition in ovarian cancer cells. Oncotarget 8, 16951–16963.

Yu, G., Wang, L.G., Han, Y., and He, Q.Y. (2012). clusterProfiler: an R package for comparing biological themes among gene clusters. OMICS 16, 284–287.

Zhang, X., Yalcin, S., Lee, D.F., Yeh, T.Y., Lee, S.M., Su, J., Mungamuri, S.K., Rimmele, P., Kennedy, M., Sellers, R., et al. (2011). FOXO1 is an essential regulator of pluripotency in human embryonic stem cells. Nat Cell Biol 13, 1092–1099.

Zhang, Y., Ma, Y., Huang, Y., Zhang, Y., Jiang, Q., Zhou, M., and Su, J. (2020). Benchmarking algorithms for pathway activity transformation of single-cell RNA-seq data. Computational and structural biotechnology journal 18, 2953–2961.

Zheng, Y., Xue, X., Shao, Y., Wang, S., Esfahani, S.N., Li, Z., Muncie, J.M., Lakins, J.N., Weaver, V.M., Gumucio, D.L., et al. (2019). Controlled modelling of human epiblast and amnion development using stem cells. Nature 573, 421–425.

Zhou, Y., Zhou, B., Pache, L., Chang, M., Khodabakhshi, A.H., Tanaseichuk, O., Benner, C., and Chanda, S.K. (2019). Metascape provides a biologist-oriented resource for the analysis of systems-level datasets. Nature communications 10, 1523.

